# LINE1 promotes nuclear compartmentalization to repress the 8-cell state in embryonic stem cells

**DOI:** 10.1101/2024.09.22.614332

**Authors:** Juan Zhang, Lamisa Ataei, Liang Wu, Kirti Mittal, Linh Huynh, Shahil Sarajideen, Abdul Mazid, David P. Cook, Daniel Trcka, Kevin Tse, Jeffrey L. Wrana, Michael M. Hoffman, Miguel A. Esteban, Miguel Ramalho-Santos

## Abstract

The family of LINE1 transposable elements underwent a massive expansion in mammalian genomes. While traditionally viewed as a mutagenic selfish element, recent studies point to roles for LINE1 in early mouse development, T cell quiescence and neurogenesis. Here we show that human LINE1 RNA is essential for self-renewal and identity of human embryonic stem cells (hESCs). Silencing of LINE1 using either antisense oligonucleotides or CRISPR interference in naïve hESCs leads to a strong induction of 8C-like cells (8CLCs). We found that genes derepressed upon LINE1 KD are not uniformly distributed across the genome, with an enrichment for chromosome 19, which includes key markers of the 8C state such as *TPRX1*. Silencing of *TPRX1*, but not other putative 8C regulators *p53* or *H3.XY*, suppresses the induction of the 8C program in LINE1 KD hESCs. We found that LINE1 RNA is preferentially localized to the lamina and periphery of the nucleolus in hESCs. Sequencing of Lamina-Associated Domains (LADs) and Nucleolus-Associated Domains (NADs) reveals a preferential association of chromosome 19 with NADs in hESCs. However, 8CLCs have a distinct nucleolar morphology and a lower association of chromosome 19 and *TPRX1* loci with the nucleolus relative to naïve and primed hESCs, suggesting a role for nucleolar dynamics in the 8CLC-hESC transition. In agreement, LINE1 KD leads to disruption of nucleolar architecture with signs of nucleolar stress. Independent perturbations of the nucleolus induce the 8C program in hESCs. Genes induced by LINE1 KD are enriched for targets of Polycomb Repressive Complex (PRC2), and inhibition of PRC2 leads to a strong induction of 8C genes. Our results indicate that LINE1 coordinates nuclear compartmentalization and chromatin-mediated gene repression to prevent developmental reversion of hESCs.

**Highlights:**

- Knockdown of LINE1 induces TPRX1-dependent emergence of 8C-like cells in hESCs.
- Genes de-repressed upon LINE1 KD are enriched for Chr 19 and PRC2 targets.
- 8CLCs display dissociation of Chromosome 19 from the nucleolus.
- Disruption of the nucleolus or inhibition of PRC2 strongly induce the 8C program.

## Introduction

Understanding early human development is of fundamental interest and may provide new applications in Reproductive Health and Regenerative Medicine. However, work in this field is hindered by ethical questions and the low availability of early human embryos for research. Human embryonic stem cells (hESCs) provide a powerful platform to model human development. Depending on culture conditions, hESCs can capture the naïve state of the pre-implantation blastocyst or the primed state of the post-implantation embryo. These two states can be interconverted as well as used to differentiate cells towards specific post-gastrulation lineages. Reversion of naïve hESCs to earlier developmental stages has been more difficult to achieve. Or particular interest is the 8-cell (8C) state, the stage at which zygotic genome activation occurs^1^. Recently, several groups reported that 8C-like cells (8CLCs), which express a distinct transcriptional program similar to the 8C embryo, exist at low frequency in cultures of naïve hESCs^2–5^. These modified hESC culture conditions represent an unprecedented opportunity to model and study human pre-implantation development. In support of this notion, the transcription factor *TPRX1* was shown to be essential both for development of human 8-cell embryos ex vivo^6^ and for generation of 8CLCs from hESCs^3^. However, it remains unclear what molecular factors prevent hESCs from reverting to the 8CLC state.

Studies in mouse have revealed several repressors of the 2-cell (2C) state^7–10^, the equivalent to the human 8C state. We and others have shown that the expression of the largest family of mammalian transposable elements (TEs), long interspersed nuclear element 1 (LINE1), is essential for mouse embryonic stem cell (mESC) self-renewal and pre-implantation development^7,11^. We found that mouse LINE1 acts as a nuclear non-coding RNA that interacts with Nucleolin (NCL) at chromatin to repress the mouse 2C program^7,12^ (reviewed in^13^). This study left several important questions unanswered, notably how LINE1 RNA might regulate chromatin organization to maintain the mESC state. Moreover, these findings raised the question of whether and how LINE1 could play a conserved role in between mouse and human. Unlike unique protein-coding genes, TEs are typically not conserved in different mammalian genomes, but rather different TE sub-families are lost and gained independently in different lineages over evolution^14–16^. This mobility and replacement of TEs makes them ideal drivers of genome evolution, as has been well documented in mammals^15,17–19^, but makes it less straightforward to envision TEs regulating conserved processes such as development or physiology across species.

In this study, we set out to determine the function of LINE1 RNA in hESCs. Our findings support a model whereby LINE1 RNA is essential for self-renewal and identity of hESCs, by preventing reversion to the 8CLC state in a manner dependent on *TPRX1*. In parallel, our data uncover key novel regulatory features of the human 8CLC state, including a role for nucleolar architecture and a unique localization of chromosome 19 and a repressive function for PRC2.

## Results

### LINE1 RNA regulates self-renewal and the transcriptional program of hESCs

We began by analyzing the expression of LINE1 subfamilies^20^ during human preimplantation development^21–23^. The expression of recently evolved primate LINE1 subfamilies is detected in oocytes and rises further after fertilization, with a peak at the inner cell mass (ICM) of the blastocyst (Figures 1A, S1A-S1B and S1D). In contrast, older mammalian LINE1 subfamilies do not display a clear pattern of expression during early human development (Figures S1C and S1E). We then determined the expression of primate LINE1 in cultures of hESCs that recapitulate pre- and post-implantation human development (Figures S1F-S1H)^3,24–26^. We found that LINE1 RNA is expressed in hESCs in all culture conditions tested and is primarily localized to the nucleus, with some cytoplasmic foci also observed (Figure S1H).

**Figure 1.**
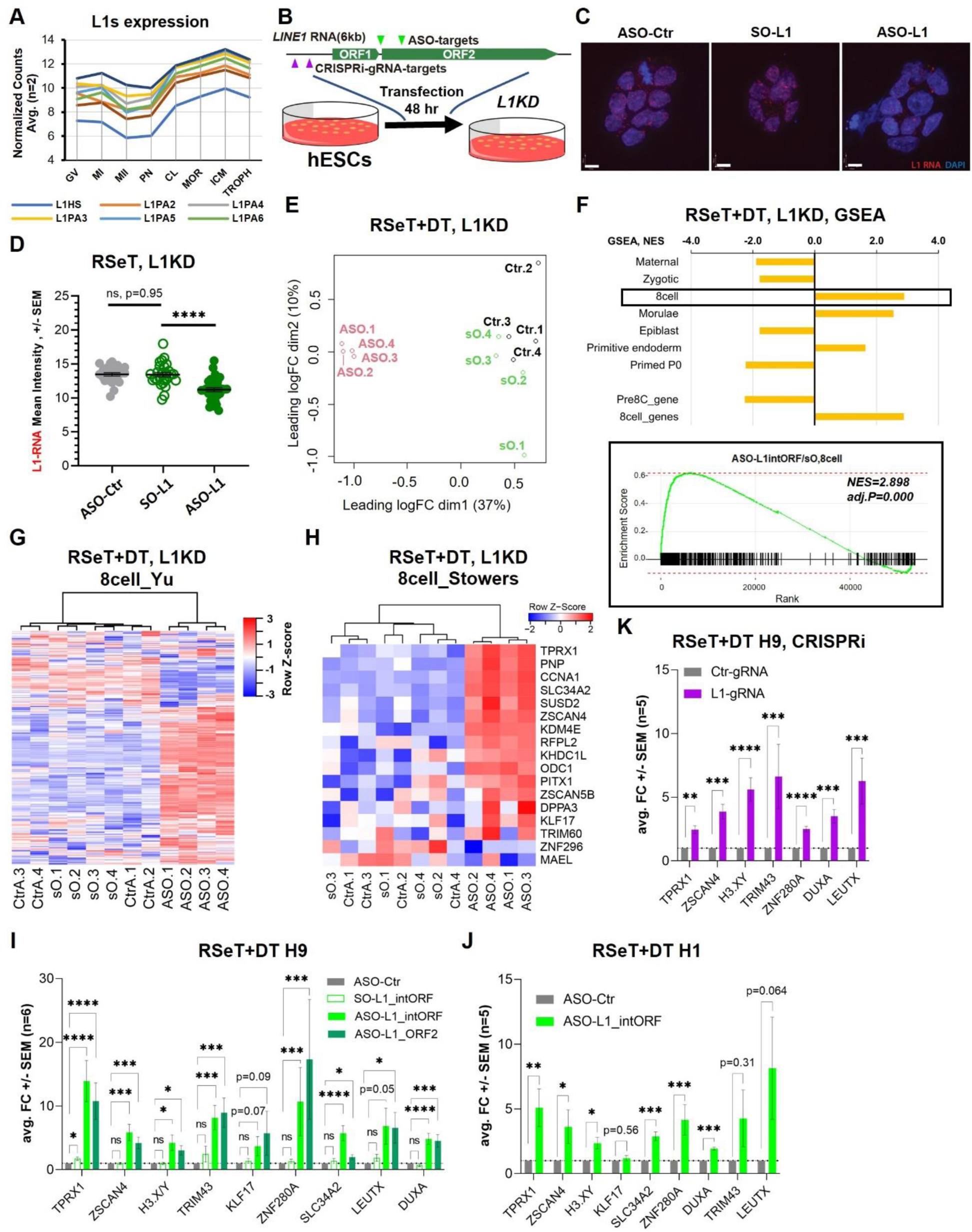
LINE1 RNA is a barrier to the 8C program in hESCs. (A) Averaged normalized counts of n=2 independent RNA-seq samples^21^ showing induction of human- and primate-specific LINE1 subfamilies (L1HS and L1PA2 to L1PA8) during human cleavage embryonic stages. GV, germinal vesicle; MI, metaphase I; MII, metaphase II; PN, pronuclear stage; CL, cleavage; MOR, morula; ICM, inner cell mass; TROPH, trophectoderm. (B) Schematic of LINE1 RNA knockdown experiments in hESCs (see Figure S1F for hESC culture conditions used in this study). ASOs targets (green arrows) at the inter-ORF and ORF2 sites, and CRISPRi gRNA targets (purple arrows) at the 5’-UTR of full-length LINE1 elements are indicated. Cells were collected after 48 hours of transfection. (C) LINE1 RNA FISH in RSeT hESCs showing predominantly nuclear localization of LINE1 RNA and efficient knockdown in ASO-L1 transfected cells compared to ASO-Ctr and SO-L1 controls after 48 hours of transfection. Representative of two independent experiments. Scale bar, 10 µm. (D) Quantification of LINE1 RNA-FISH signal intensity in (C). LINE1 intensity in the ASO-L1 transfected cells is significantly reduced compared to the controls. Representative of three independent experiments. See Extended Data 2a-b for validation of LINE1 KD in RSeT+DT and primed hESCs. Mann-Whitney test. ns = p > 0.05, ** = p < 0.01. (E) Multidimensional scaling (MDS) plot of genes across all samples, showing that ASO-L1 KD RSeT+DT hESCs have distinct gene expression profiles from SO-L1 and ASO-Ctr samples. (F) Gene Set Enrichment Analysis of the transcriptional profile of ASO-L1 KD RSeT+DT hESCs compared to different stages of peri-implantation development and 8C/pre-8C gene sets^4,22^. LINE1 KD hESCs display a significant enrichment for the 8-cell (plotted in the lower panel) and morulae programs. NES, Normalized Enrichment Score. (G and H) Heatmap showing induction of diagnostic gene sets of the 8C stage in vivo from X. Yu et al.^4^ (G) and Taubenschmid-Stowers et al.^2^ (H) in RSeT+DT hESCs upon ASO-L1KD compared to ASO-Ctr and SO-L1 controls. (I) qRT-PCR validation of upregulation of 8C genes in LINE1 KD RSeT+DT H9 hESCs with ASOs targeting the inter-ORF and ORF2 sites, respectively. Data are mean ± SEM, n = 6 biological replicates. Ratio paired Student’s t-tests. ns = p > 0.05, * = p < 0.05, ** = p < 0.01, *** = p< 0.001, **** = p < 0.0001. (J) qRT-PCR validation of upregulation of 8C genes in LINE1 KD RSeT+DT H1 hESCs with ASOs targeting the inter-ORF site. Data are mean ± SEM, n = 5 biological replicates. Ratio paired Student’s t-tests. ns = p > 0.05, * = p < 0.05, ** = p < 0.01, *** = p< 0.001. (K) qRT-PCR validation of upregulation of 8C genes in LINE1 KD RSeT+DT H9 hESCs with CRISPRi targeting LINE1. Data are mean ± SEM, n = 5 biological replicates. Ratio paired Student’s t-tests. ** = p < 0.01, *** = p< 0.001, **** = p < 0.0001.

To investigate the function of LINE1 in hESCs, we designed an antisense oligo (ASO) targeting a sequence conserved in the interORF region of primate and human-specific LINE1 families (ASO-L1) (Figure 1B; Table S1). We found that ASO-L1 induces efficient knockdown of LINE1 RNA (L1KD) relative to a control non-targeting ASO (ASO-Ctr) or a sense version of the ASO-L1 (SO-L1) across different hESC culture conditions (Figures 1C-1D and S2A-S2C). Regardless of culture conditions, knockdown of LINE1 leads to loss of self-renewal of hESCs (Figures S3A-S3F), consistent with results observed in mESCs^7^.

To determine the impact of LINE1 KD on the transcriptome of hESCs, we carried out bulk RNA-sequencing (RNA-seq). Three different culture conditions of karyotypically normal (Figures S4A-S4B) H9 hESCs were tested, which capture the pluripotency continuum between naïve and primed states (Figure S1F): mTeSR medium, to maintain a primed state^26^, RSeT medium, to maintain an intermediate naïve-like state^24^ (a RT-PCR validation in Figure S1G), and RSeT+DT, where RSeT medium is supplemented for 48 hours with 10nM DZNep (3-Deazaneplanocin A, “D”), an S-adenosyl-L-homocysteine hydrolase, and 5nM TSA (Trichostatin A, “T”), a histone deacetylase inhibitor. These low doses of DT were recently shown to promote a more naïve state permissive for the emergence of 8CLCs at low frequencies^3^ (Figure S1F). These conditions have the advantages of relying on standardized commercial base media (mTeSR and RSeT) and avoiding prolonged cultures in DT, which we found can lead to increased expression of 8C markers and cell death over time (data not shown). Across all three hESC culture conditions tested, LINE1 KD induces reproducible changes in gene expression relative to controls, as determined by principal component analysis and hierarchical clustering (Figures 1E, 1G, S5A-S5E and S5G-S5H).

We next performed Gene Set Enrichment Analysis (GSEA) to compare the transcriptional changes induced by LINE1 KD to gene sets that define the stages of human preimplantation embryos from published data^4,22^ (Figure S5M; Table S2). This analysis revealed that LINE1 KD induces transcriptional signatures of the 8C and morulae stages, which are not normally expressed in hESCs (Figures 1F and S5J-S5K). Hierarchical clustering confirmed that the 8C gene expression program^4^ is induced upon LINE1 KD in hESCs (Figures 1G and S5G-S5H). While derepression of the 8C signature is detected upon LINE1 KD across all hESC culture conditions tested, this effect is more prominent in cells of a more naïve state (RSeT+DT > RSeT > primed) (Figures 1F and S5J-S5L). Moreover, a short diagnostic set of 8C markers^2^ is robustly induced by LINE1 KD in RSeT+DT (Figures 1H-1I, S5C and S5F) but much less so in RSeT (Figures S5D and S5I), suggesting that a permissive chromatin environment induced by DT facilitates 8C gene derepression upon LINE1 KD (Figures S1F and S5L). Importantly, the induction of diagnostic 8C markers upon LINE1 KD was validated using an independent ASO targeting a conserved region in ORF2 (ASO-L1_ORF2), or using a different, XY H1 hESC line (Figures 1I-1J). Moreover, we corroborated our findings using a CRISPR interference (CRISPRi) system in hESCs generated via sleeping beauty transposition^27^. Transfection of gRNAs to target a stably expressed dCas9-KRAB repressor protein to the LINE1 promoter region^28,29^ (Figure S2D; Table S1) leads to knockdown of LINE1 RNA (Figure S2E) and induction of 8C marker genes (Figure 1K). Taken together, these results indicate that knockdown of LINE1 RNA via independent methods consistently leads to the derepression of the 8C program in hESCs.

### LINE1 KD induces the generation of 8CLCs

8CLCs are expected to be a rare subset of cells in naïve hESC cultures^2–5^. To determine whether the upregulation of the 8C program upon LINE1 KD arises from the induction of 8CLCs, we carried out 10X Genomics single-cell RNA-seq (scRNA-seq) on hESCs transfected with SO-L1 (control) or ASO-L1 (LINE1 KD). A total of 5471 cells in control and 7629 cells in LINE1 KD were sequenced and passed quality control, respectively (Figures 2A and S6A-S6B). Uniform manifold approximation and projection (UMAP) shows that a population of cells expresses 8C gene signatures^22,3^, including key 8C markers such as *TPRX1*, *ZSCAN4*, *DPPA3*, *ZNF280A* and *SLC34A2* (encircled in Figure 2A and cluster 15 in Figure S6B). This cluster of cells co-expresses TEs previously reported as markers of the 8C stage in vivo and 8CLCs in vitro, such as *MLT2A1/2* and *LTR7B*^2,3,5^ (Figure 2B). As expected from 8CLCs, this cluster also expresses lower levels of pluripotency markers such as *POU5F1*, *SOX2* and *NANOG* (Figures 2C and S6C-S6D). Some 8C markers (such as *TPRX1*, *DPPA3*, and *CCNA1)* are consistently detected across all cells of this cluster, while others (*DUXA*, *KLF17*, and *ZSCAN4*) are detected in only some of them (Figures 2A, and 2C). These differences may reflect cellular heterogeneity within the 8CLC population or limitations of sequencing depth. Nevertheless, it is clear that the 8CLC cluster of cells is essentially only observed upon LINE1 KD: while only 0.07% (4/5471) of control hESCs belong to the 8CLC cluster, LINE1 KD induces a ∼65-fold enrichment, to 4.95% (378/7629) of the population.

**Figure 2.**
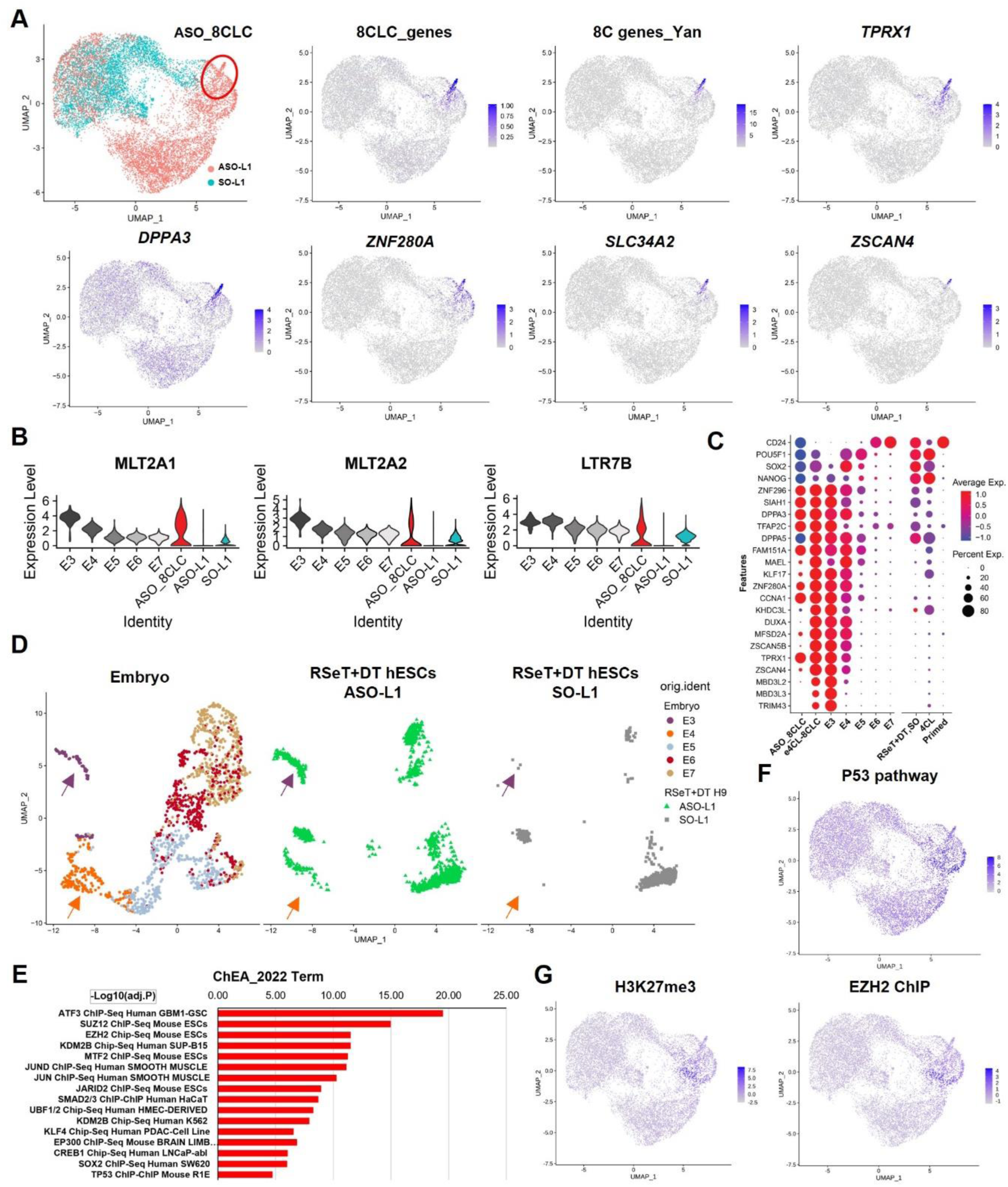
LINE1 KD induces the generation of 8CLCs. (A) UMAP visualization of scRNA-seq data from LINE1 KD and control RSeT+DT hESCs. The top left panel displays ASO-L1 KD and SO-L1 control cells as red and blue dots, respectively. The circled population (cluster 15 in Figure S6B) represents LINE1 KD-induced 8CLCs (ASO_8CLCs) expressing 8C gene sets annotated from 8CLCs^3^ and 8C embryos^22^; individual 8C marker genes are also indicated. (B) Violin plot showing the log-normalized expression of representative 8C-enriched TEs (*MLT2A1*, *MLT2A2* and *LTR7B*) in human E3 to E7 stages, ASO-L1 KD-induced 8CLCs (ASO_8CLCs, circled in (A), upper left panel), and total hESCs transfected with ASO-L1 or SO-L1 (red and blue dots in (A), upper left panel). (C) Bubble plot representing the frequency of expression and average expression of representative pluripotency and totipotency genes in early human embryonic stages^30^, ASO-L1 KD-induced 8CLCs (ASO_8CLCs), e4CL induced 8CLCs (e4CL_8CLCs)^3^, RSeT+DT control hESCs (SO-L1 transfected), 4CL hESCs and primed hESCs. (D) UMAP visualization of scRNA-seq data from early human embryos (E3 to E7)^30^. To the right, data from ASO-L1 or SO-L1 transfected hESCs are projected onto the in vivo data. Purple and orange arrows point to corresponding E3 and E4 cell clusters, respectively. (E) Top chromatin-bound factors enriched at genes upregulated upon L1KD in RSeT+DT hESCs (see Methods). (F) UMAP view of expression levels of genes of the p53 pathway. (G) UMAP view of expression levels of H3K27me3-marked genes (left) and targets of EZH2 (right) in hESCs.

To directly assess the similarity of LINE1 KD hESCs to embryonic cells in vivo, we mapped our scRNA-seq data onto published scRNA-seq of early human embryos^30^. This analysis revealed that coherent groups of cells in the LINE1 KD dataset, but not the control dataset, are projected onto the cell clusters of embryonic day (E) 3 (8C) and E4 (morula) embryos (Figure 2D).

In addition to the induction of a distinct 8CLC cluster, there are other transcriptional changes induced by LINE1 KD that potentially reflect a more general transcriptional deregulation. We analyzed the bulk RNA-seq dataset of LINE1 KD hESCs for enriched molecular signatures (Figures 1E-1H, S5C and S5F) and projected them onto the scRNA-seq data. This analysis revealed that cell clusters induced by LINE1 KD are enriched for genes upregulated upon silencing of the pluripotency factors *POU5F1* or *SOX2* (Figures S6E-S6F), signatures of stress such as the p53 pathway (Figures 2E-2F and S6G), and targets of the Polycomb Repressive Complex 2 (PRC2), which deposits H3K27me3 (Figures 2E, 2G and S6H-S6I, see below).

To independently assess the emergence of 8CLCs in hESC cultures, we used immunofluorescence (IF) to assess the expression of TPRX1 protein, a diagnostic marker of 8CLCs^2,3^. In H9 hESCs, LINE1 KD induces an increase in TPRX1+ cells from ∼0.8% to ∼3.6% (Figures 3A-3B); in H1 hESCs, LINE1 KD induces an increase from ∼0.2% to ∼4.2% (Figures S7A-S7B). These percentages roughly parallel those derived from the scRNA-seq data. Taken together, these results indicate LINE1 KD induces the emergence of 8CLCs in hESCs.

**Figure 3.**
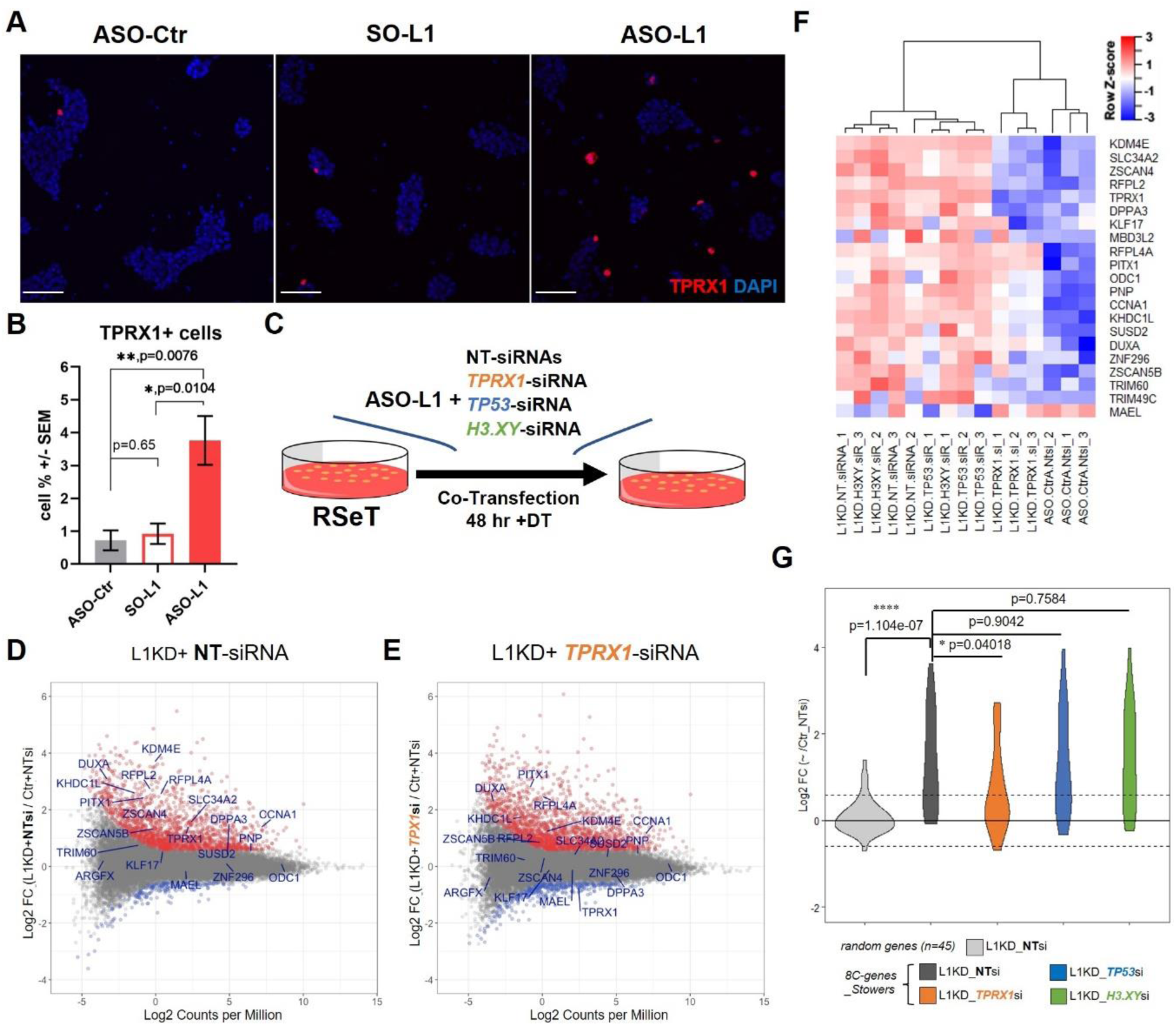
The induction of the 8C program upon LINE1 KD is mediated by *TPRX1*. (A) Immunostaining showing an increase in TPRX1-positive cells in RSeT+DT H9 hESCs upon ASO-L1KD compared to controls. Representative of three independent experiments. Scale, 100 µm. (B) Quantification of (A) for percentage of TPRX1+ cells in each indicated sample. 1000∼1500 cells per sample from six randomly scanned images of two independent experiments are quantified. Data are mean ± SEM. Welch’s t-test. * = p < 0.05, ** = p < 0.01. Representative of three independent experiments. (C) Schematic of LINE1 RNA knockdown, co-transfected with siRNAs targeting *TPRX1*, *TP53*, and *H3.XY*, compared with a control non-targeting (NT) siRNA. Cells were collected after 48 hours of transfection (D and E) MA plot showing log2 fold change in gene expression following (E) L1 KD+NT-siRNA and (F) L1 KD+*TPRX1*-siRNA compared to control transfected cells (ASO-Ctr+NT-siRNA). Red or blue highlight genes of adj.P-value < 0.05 and FC > 1.5 or < −1.5, respectively. 8C marker genes from Taubenschmid-Stowers et al.^2^ are labeled in dark blue. (F) Heatmap showing a suppression in the induction of 8C marker genes when LINE1 KD is combined with *TPRX1* RNAi but not with *TP53* or *H3.XY* RNAi. (G) Violin plot of the same data as in (F). *TPRX1* RNAi, but not *TP53* or *H3.XY* RNAi, significantly suppresses the induction of 8C marker genes that is observed upon LINE1 KD, relative to an random gene set of equal size. Wilcoxon test, * = p < 0.05, **** = p < 0.0001.

### The induction of the 8C program upon LINE1 KD is mediated by *TPRX1*

We next sought to identify genes that may mediate the induction of the 8C program upon LINE1 KD. Several studies have suggested that the transcription factor *DUX4* is a master regulator of the human 8C program^21,31,32^, similar to the role of its homolog *Dux* in mouse^21,33,34^. In mESCs, the induction of the 2C program upon LINE1 KD is *Dux*-dependent^7^. However, we found that *DUX4* and *DUX4*-like genes are essentially undetectable in our RNA-seq data and are not induced by LINE1 KD (Figure S7C). This is in contrast to *TPRX1* (see above, Figures 1, 2, and S7D), a known regulator of human 8CLCs in vitro^3^ and 8C development in vivo^6^. In addition to *TPRX1*, we considered a potential role for *H3.XY*, a histone variant that is a marker of the 8C state^2^ induced by LINE1 KD (Figures 1I-1K) which mediates the induction of *DUX4* target genes in human muscle cells^35^. Finally, we considered *TP53*, as it has been linked to stress-induced activation of 2C/8C genes in mouse and human cells^36,37^ and p53 pathway genes are induced by LINE1 KD (Figures 2E-2F).

To test the potential regulatory roles of these candidate genes, we carried out ASO-mediated LINE1 KD with or without simultaneous KD of *TPRX1*, *H3.XY* or *TP53* using siRNAs (Figure 3C). We validated that *TPRX1, H3.XY* and *TP53* levels are knocked down by their corresponding siRNAs (Figures S8A-S8D). RNA-seq revealed that KD of *TPRX1*, but not *H3.XY* or *TP53,* partially suppresses the induction of the 8C program upon LINE1 KD (Figures 3D-3G and S8E-S8G). In particular, the expression of key diagnostic markers of 8CLCs^2^, including *KDM4E*, *SLC34A2*, *ZSCAN4*, *RFPL2*, *DPPA3* and *KLF17* (apart from *TPRX1* itself), is reverted towards control levels in LINE1 KD cells only in the case of *TPRX1* RNAi (Figures 3E-3F). Note that there are transcriptional changes induced by LINE1 KD not related to 8C program that are not rescued by *TPRX1* RNAi (Figures 3D-3E). We cannot exclude the possibility that RNAi did not lead to sufficient KD of *H3.XY* or *TP53*. Moreover, there are likely other mediators of the transition to 8CLCs, in addition to *TPRX1*. Nevertheless, our findings indicate that the induction of the 8C program upon LINE1 KD is partially mediated by derepression of *TPRX1*.

### Lamina- and nucleolus-associated domains are enriched for LINE1 and reveal a dynamic localization of 8C loci

In mouse 2CLCs, the activation of *Dux* coincides with its relocalization from the nucleolar periphery to the nucleoplasm^7,38,39^, although the underlying mechanisms remain poorly understood. We therefore sought to understand the dynamics of nuclear compartmentalization, including the localization of LINE1 and 8C genes, in hESCs. We carried out sequencing of NADs (using nucleolar fractionation-seq^40^) and LADs (using LAMB1-DamID-seq^41,42^) in hESCs (Figure 4A). Both of these compartments are associated with heterochromatin-mediated gene repression^43,44^. While NADs have not previously been sequenced in hESCs, our LAD-seq data are highly consistent with a previous report^45^. We found that human LINE1 loci are enriched in NADs and LADs in hESCs (Figures S9A-S9B), in agreement with data in mESCs^12,46^. Interestingly, we found that LINE1 RNA accumulates in nuclear foci that are frequently located at the vicinity of the nucleolus and lamina, in contrast to a lower density in the intervening nucleoplasm (Figures 4B and S9C-S9D). Moreover, cross-linking and RNA immunoprecipitation (CLIP)-qRT-PCR revealed an interaction between Nucleolin (NCL) and Laminin B1 (LMNB1) proteins with LINE1 RNA (Figure S9E). In addition, we found that LADs are essentially constant between naïve-like and primed hESCs, while NADs are less abundant in naïve-like hESCs and expand in signal and size in primed hESCs (e.g., see Figures S12A). These results suggest that the nucleolar compartment may be particularly dynamic during early human development (see below).

**Figure 4.**
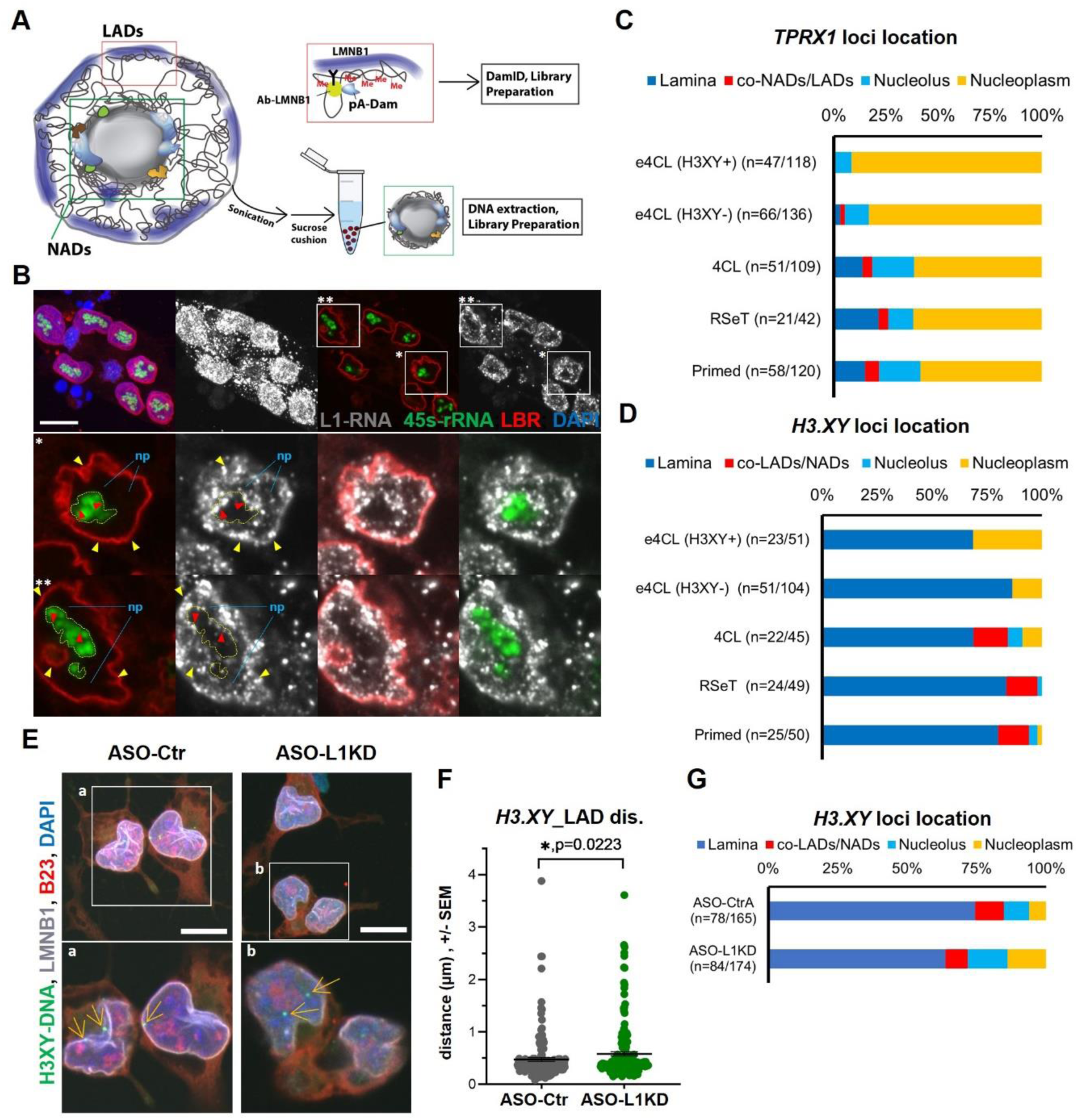
Lamina- and nucleolus-associated domains are enriched for LINE1 and reveal a dynamic localization of 8C loci. (A) Schematic of LAD-seq and NAD-seq in hESCs. See Methods for details. (B) Representative image of LINE1 and 45s-rRNA RNA-FISH and co-IF staining for LBR antibody in RSeT+DT hESCs, showing that LINE1 RNA are enriched in the vicinity of nucleolar and laminar domains. The max projected image (two top left panels) displays high expression of LINE1 in the nucleus. A representative single-plane is shown in the two top right panels, with boxed areas (**\*** and **\*\***) enlarged in the middle and bottom panels, with color channels separated. Yellow dotted line encircles the nucleolus area. Yellow and red arrows point to LINE1 RNA foci-enriched spots at the nucleolar and laminar domains, respectively. Noted the nucleoplasm (np) areas are sparse in LINE1 foci. Representative of at least three independent experiments. Scale bar, 10 µm. (C) Distribution of the location of *TPRX1* loci across the nuclear compartments listed. See also Figures S10. n = cells /*TPRX1* loci quantified. (D) Distribution of the location of *H3.XY* loci across the nuclear compartments listed. See also Figures S11. n = cells /*H3.XY* loci quantified. (E) Representative image of *H3.XY* DNA-FISH and and co-IF staining for B23 and LMNB1 in control and LINE1 KD RSeT+DT hESCs. *H3.XY* loci localize further away from the lamina in LINE1 KD cells. The boxed area in the max projected images (top panels) is enlarged and displayed as representative z-stack (Z) images (bottom panels). The yellow arrows point to *H3.XY* loci in the nucleoplasm. Representative of three independent experiments. Scale bar, 10 µm. (F) Quantification of data from *H3.XY* DNA-FISH and co-IF staining (shown in E), plotting the distance of *H3.XY* loci to the lamina domain. If the distance is < 0.5 µm, it is defined as within LADs. Number of cells quantified in each group is indicated in (G). Mann-Whitney test. * = p < 0.05. Data are from three independent experiments. (G) Distribution of the location of *H3.XY* loci across the nuclear compartments listed in control and LINE1 KD RSeT+DT hESCs based on the distance quantification in F. n=cells /*H3.XY* loci quantified. Data are from three independent experiments.

The NAD and LAD data further predicted an enrichment of 8C gene loci like *TPRX1* and *H3.XY* in the nucleolus or the lamina (Figure S10A and S11A). By DNA-FISH, we found that *TPRX1* loci are localized in the nucleoplasm in 8CLCs, but display increasing association with the nuclear lamina and the nucleolar periphery with developmental progression of hESCs (Figures 4C and S10B-S10D; Videos S3-S6). In contrast, *H3.XY* (Chr. 5, *H3.Y1* and *H3.Y2*, see Figure S11A) is located at the nuclear lamina across all hESC states, with only ∼20% of *H3.XY* loci relocated to the nucleoplasm in 8CLCs (Figures 4D and S11B-S11C). We find *DUX4* to be mostly localized to the nuclear lamina in naïve hESCs (Figures S11D-S11E), which stands in contrast to the nucleolar association of its homolog *Dux* in mESCs^7,39^. Taken together, the LAD-seq and NAD-seq data, as well as the imaging data for key loci, reveal a progressive association of 8C genes with both the nucleolar periphery and the nuclear lamina, from 8CLCs to more developmentally advanced stages of hESCs.

We next explored whether LINE1 KD impacts the 3D nuclear localization of specific 8C loci. Interestingly, we found that in LINE1 KD hESCs a lower proportion of *H3.XY* loci are located at the lamina (∼72% vs ∼85% in control cells), and their overall distance from the lamina increases (Figure 4E-4G). These results suggest that LINE1 RNA promotes repression of *H3.XY* genes in part by contributing to tethering of their loci to the nuclear lamina.

In contrast, we detected no significant change in the localization of *TPRX1* loci upon LINE1 KD (Figure S10E-S10F). The observation that only a minor percentage of *TPRX1* loci are located near the lamina or the nucleolus in naïve and primed hESCs (Figure 4C) complicates a quantification of the impact of LINE1 KD. Moreover, the fact that *TPRX1* is repressed in naïve and primed hESCs despite its predominant nucleolplasmic localization indicates that mechanisms other than 3D nuclear positioning may contribute to *TPRX1* silencing in hESCs.

### Chromosome 19 is enriched for genes induced upon LINE1 KD and preferentially associates with the nucleolus

We noticed that *TPRX1*, as well as several other key markers of the 8C state (e.g., *TPRX2, ZSCAN4, DUXA* or *LEUTX*) are all located on Chr. 19 (Figure 5A). We therefore analyzed the chromosomal distribution of genes differentially expressed in LINE1 KD hESCs. Intriguingly, Chr. 19 displays an enrichment for genes upregulated upon LINE1 KD (Figure 5B), across all hESC states tested (RSeT+DT, RSeT and Primed). Moreover, analysis of the LAD/NAD-seq data revealed that, while most chromosomes display enrichments for both compartments, Chrs. 19 and 22 stand out as being enriched for NADs but not LADs in both naïve and primed hESCs (Figures 5C and S12B). This observation is supported by data in differentiated cells showing that Chr.19 occupies an internal nuclear territory closely associated with the nucleolus^47,48^.

**Figure 5.**
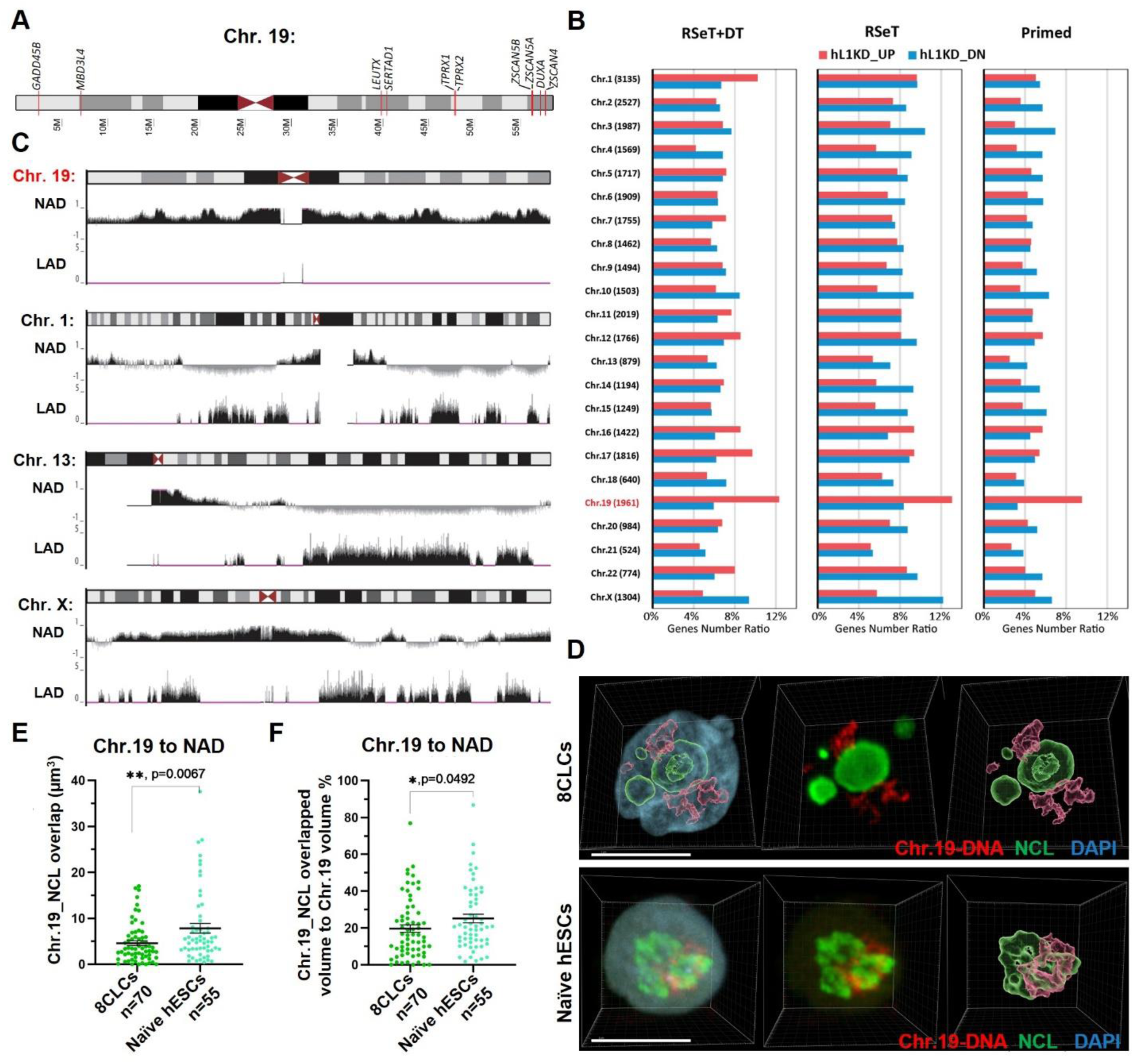
Chromosome 19 is enriched for genes induced upon LINE1 KD and preferentially associates with the nucleolus. (A) Ideogram of chromosome 19 with the location of several 8C genes indicated. (B) Distribution of protein-coding genes upregulated (red bars) or downregulated (blue bars) upon LINE1 KD as percentage of total number of genes in each chromosome. Chr. 19 shows a bias towards genes upregulated vs downregulated in comparison to other chromosomes, in all hESC culture condition used. (C) Representative genome browser views of LAD and NAD enrichment signatures of Chr. 19, Chr. 1, Chr. 13, and Chr. X. Chr. 19 displays a distinct enrichment of NAD but not LAD throughout the chromosome. (D) Example still images from videos (Videos S1 and S2) of Chr. 19 DNA-FISH and co-IF staining for NCL antibody, showing that Chr.19 is less associated with the nucleolus in H3,XY+ 8CLCs than in naïve hESCs, cultured in e4CL and 4CL^3^, respectively. 3D reconstructed fluorescence and surface data are shown, see Methods for details. Representative of two independent experiments. Scale bar, 20 µm. (E and F) Quantification of Chr.19 DNA-FISH and NCL IF co-staining (as shown in D) indicates a significantly higher overlap between these two territories in naïve hESCs than in 8CLCs (whether H3.XY+ or H3.XY-). Data are from two replicated independent experiments. Number of cells quantified in each group is indicated. Mann-Whitney test. * = p < 0.05, ** = p < 0.01.

The results above prompted us to explore the 3D nuclear positioning of Chr. 19 in 8CLCs vs hESCs. We carried out chromosome painting in enhanced 4CL medium with high levels of DT (“e4CL”), which was recently shown to enrich for 8CLCs, compared to naïve cells in basal 4CL medium^3^. The results reveal a remarkably distinct association between Chr. 19 and the nucleolus: while in naïve cells Chr. 19 is intimately associated with the nucleolus, as would be predicted from the NAD data (Figures 5C and S12), in 8CLCs there is a significantly higher separation between the two (Figures 5D-5F; Videos S1 and S2). This differential co-localization seems to be in part related to a much rounder morphology of the nucleolus in 8CLCs. Thus, 8CLCs appear to be the exception to the rule of an intimate association between Chr. 19 and the nucleolus across cell types tested. The molecular mechanisms that underlie the peri-nucleolar localization of Chr. 19, and that presumably are not yet in place in 8CLCs, remain unknown (see Discussion). Nevertheless, these results led us to focus our attention on the nucleolus for the remainder of this study.

### LINE1 RNA is required for maintenance of nucleolar architecture, which is critical to suppress the 8C program

As mentioned above, 8CLCs tend to display a rounder nucleolar morphology that is remarkably distinct from the irregular shape of the nucleolus typical of other hESC states and non-pluripotent cells (Figures 5D and S10B; Videos S1-S6). This nucleolar structure in 8CLCs resembles what is observed in vivo^49^. Early human pre-implantation embryos have round nucleoli up to 4-8 cell stage; from then onwards, the nucleolus becomes more irregular in shape and accumulates heterochromatin at the periphery^49^. We found that inhibition of RNA Pol I with 0.25 µM BMH in RSeT+DT hESCs induces a rounder nucleolar morphology by 8 hours (Figure S13A) and a striking up-regulation of 8C marker genes by 24 hours (Figure 6A). These results are in agreement with data showing that a functional nucleolus is required for repression of the 2C state in mESCs^38,39^. Thus, the nucleolus plays a conserved role in the repression of totipotency-associated gene expression programs in mouse and human ESCs.

**Figure 6.**
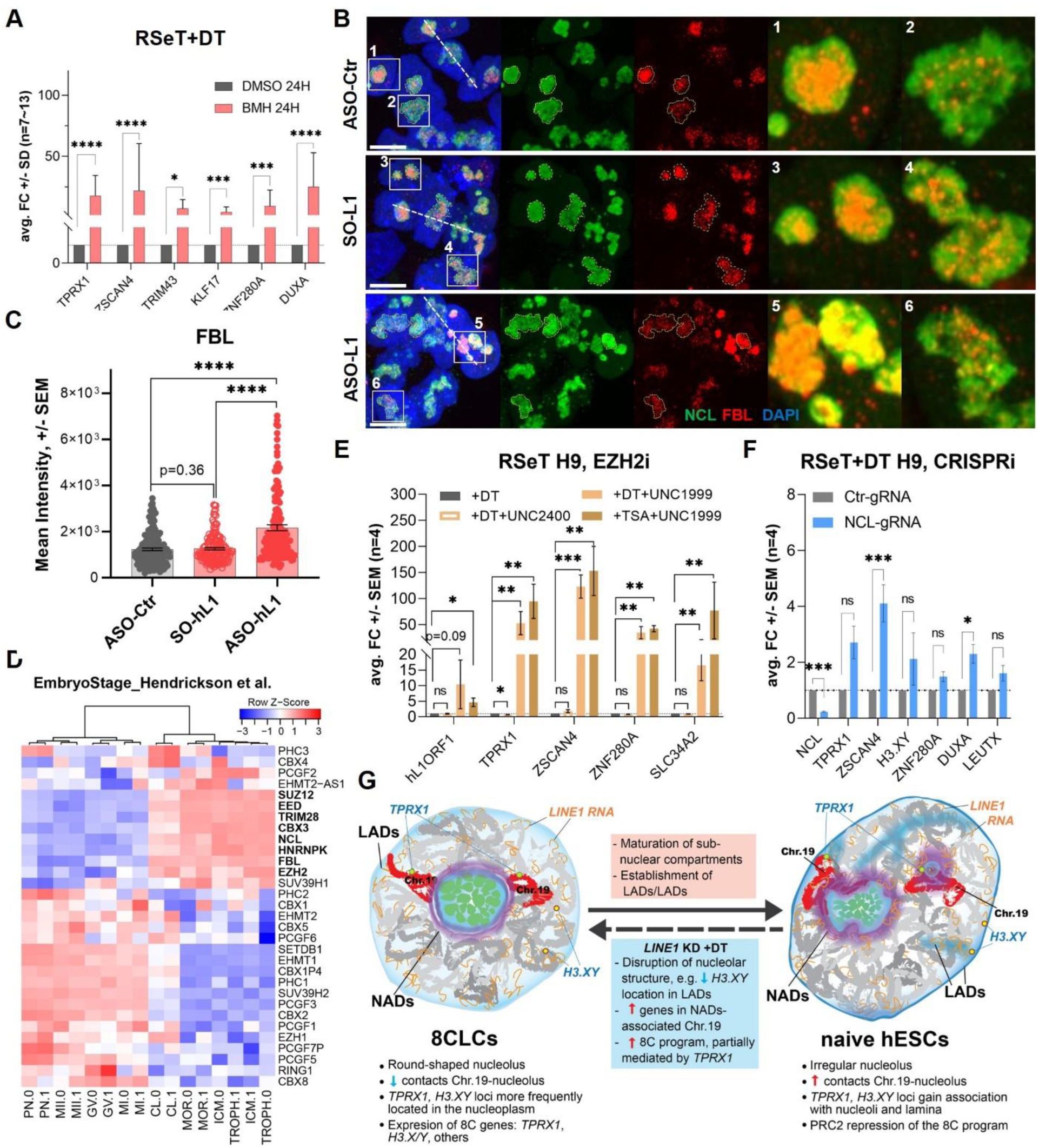
LINE1 RNA promotes maintenance of nucleolar architecture, which suppresses the 8C program. (A) qRT-PCR data showing strong upregulation of 8C genes following inhibition of RNA Pol I in hESCs using 0.25 µM BMH for 24 hours. Data are mean ± SEM, n = 7-13 biological replicates. Ratio paired Student’s t-tests. * = p < 0.05, *** = p< 0.001, **** = p < 0.0001. (B) NCL and FBL IF in control and LINE1 KD hESCs. In the LINE1 KD cells, FBL extrudes from the NCL territory. Representative of two independent experiments. Scale bar, 10 µm. (C) Quantification of the FBL IF images (as shown in B). LINE1 KD cells have significantly higher levels of FBL than control hESCs Data are from two independent experiments. Mann-Whitney test. **** = p < 0.0001. (D) Heatmap representation of the expression of select heterochromatin and nucleolus factors during early human embryonic development, profiled using bulk RNA-seq^21^. See Figure S14 for similar scRNA-seq data. Nucleolar proteins and PRC2 subunits are among the genes upregulated (bolded) at cleavage stages (CL). (E) qRT-PCR data showing upregulation of 8C genes following inhibition of EZH2 in RSeT+DT or RSeT+TSA hESCs using 2.5 µM UNC1999 for 48 hours, compared to RSeT+DT alone or RSeT+DT+UNC2400 controls. Data are mean ± SEM, n = 4 biological replicates. Ratio paired Student’s t-tests. * = p < 0.05, ** = p < 0.01, *** = p< 0.001. (F) qRT-PCR data showing upregulation of 8C genes upon knockdown of NCL by CRISPRi. Data are mean ± SEM, n = 5-7 biological replicates. Ratio paired Student’s t-tests. * = p < 0.05, ** = p < 0.01. (G) Model for the maturation of NADs and LADs, and associated 3D nuclear redistribution of Chr. 19 and key 8C loci between 8CLCs and naïve hESCs. Human LINE1 RNA promotes maintenance of nucleolar architecture and repression of the 8C program. See text for details.

We next assessed the role of LINE1 in nucleolar architecture. Normal nucleoli are organized with an interior dense fibrillar component marked by Fibrillarin (FBL), encased by an exterior granular component marked by NCL^50^. We found that LINE1 KD hESCs display a disruption of this organization and other signs of nucleolar stress^51,52^. In LINE1 KD hESCs, the FBL domain frequently extrudes onto the outside of the NCL domain, forming what are known as nucleolar caps (Figures 6B and S13B). In addition, LINE1 KD hESCs display abnormally high levels of FBL (Figure 6B-6C and S13B), which have been reported for nucleolar stress in human cells^52^. LINE1 KD leads to a disruption of nucleolar morphology than is overall distinct from the “circularization” observed upon inhibition of RNA Pol I (Figure S13A). These results indicate that LINE1 RNA is essential to globally maintaining a mature nucleolar architecture in hESC cultures, which in turn prevents a fraction of the cells from reverting to the 8CLC state.

To gain further insight into factors that might regulate exit from the 8C state, we examined the expression of various heterochromatin repressors and nucleolar proteins during early human development. Interestingly, this analysis revealed that core components of the PRC2 complex (EZH2, EED, SUZ12), as well as key nucleolar proteins (NCL and FBL), are highly induced at cleavage stages (Figures 6D and S14A-S14C), consistent with the induction of primate LINE1 subfamilies (Figures 1A, S1A-S1B and S1D). This contrasts with to PRC1 complex members or factors involved in H3K9 methylation, which are either not detected or show the oppositive pattern of expression during early human development (Figures 6D, S14A and S14D-S14G). Together with the enrichment of genes induced upon LINE1 KD for targets of PRC2 (Figures 2E, 2G and S6H-S6I), these results led us to explore a potential role for PRC2 in 8C gene regulation. Remarkably, inhibition of PRC2 using UNC1999 (but not its inactive analog UNC2400) for just 48h induces a strong up-regulation of 8C genes, even in the absence of DZNep in the culture medium (Figure 6E). In parallel, CRISPRi-mediated repression of NCL also up-regulates the expression of key 8C marker genes (Figures 6F and S13C-S13D). Taken together, these findings indicate that nucleolar maturation promoted by LINE1 RNA and NCL, possibly acting in combination with PRC2-mediated gene repression, prevents reversion of hESCs to the 8C state.

## Discussion

The regulation of the human 8C state remains very poorly understood. The transcription factor TPRX1, part of a larger family of fast-evolving homeobox transcription factors^53,54^, does not appear to play a role in mouse but is essential for the generation of 8CLCs^3^ in hESCs and development of human 8C embryos^6^. Here we show that LINE1 is an essential regulator of hESC self-renewal and identity by preventing developmental reversion to the 8CLC state. We found that LINE1 RNA is associated with the nuclear lamina and the nucleolus and is required for the maintenance of nucleolar architecture, acting as a barrier to the expression of *TPRX1* and the activation of the 8C program in hESCs (Figure 6G). In addition, our study provides new insights into chromosome 19 territory dynamics and a key organizing role for the nucleolus within the totipotency-to-pluripotency continuum of hESCs. Our analyses further point to a key role for PRC2 in repression of the 8C program. Taken together, our results indicate that LINE1, rather than being irrelevant junk DNA^55^ or a genomic parasite detrimental to cell survival^56^, coordinates 3D nuclear organization and chromatin-mediated gene repression in hESCs.

Our findings indicate that a fundamental transition at the onset of development is regulated in both mouse^7^ and human (this study) by the expression of LINE1 elements. We speculate that the massive expansion of LINE1 in therian mammals^57^ may have contributed to its role in nuclear compartmentalization and early development. Notably, the partial similarities between the roles of LINE1 in mouse and human ESCs are observed despite overall replacement of LINE1 sub-families and lack of syntheny in genomic locations of their elements across mammals^14–16,58,59^. LINE1 elements in both the mouse and human genome are located in AT-rich, gene poor areas that tend to be silenced and enriched in LAD and NAD compartments^12,46,58,59^. The AT-rich consensus sequence of the LINE1 endonuclease (5’-TTTT/AA-3’ and derivatives)^60,61^ and the selection for maintenance of LINE1 insertions in heterochromatin areas and/or against their retention in gene-rich euchromatic areas^62,63^ may contribute to preserving a role of LINE in genome compartmentalization. It will be of interest to study potential developmental roles of LINE1 and its distribution across nuclear compartments in other mammalian species.

This study points to a remarkable remodeling of the nucleolar compartment in the transition between 8CLCs and naïve hESCs. We speculate that the strong induction of key nucleolar factors at the 4-8C stage (Figures 6D and S14) may support the maturation of the nucleolus that occurs at these stages^49^. The association of LINE1 RNA with newly synthesized NCL protein may facilitate liquid-liquid phase separation^64,65^ and/or maturation of the nucleolus. In turn, the nucleolar periphery may then become a hub for recruitment of chromosomal domains to be repressed, including large parts of Chr. 19, and heterochromatin-mediated silencing of 8CLC regulators such as *TPRX1* (Figure 6G).

Our findings raise several challenging questions for future inquiry: i) what mediates the preferential association of Chr. 19 with the nucleolus in hESCs and other human cells^47,48^, and what makes this association weaker in 8CLCs? The link to LINE1 could well be indirect via a role of LINE1 in nucleolar maturation, as Chr. 19 has an overall lower frequency of LINE1 elements relative to the rest of the genome^66^; ii) How is the 3D nuclear compartmentalization of hESCs coordinated with chromatin-level repression of the 8C program? Our results indicate that PRC2 plays a key role in silencing the 8C program, and PRC2 has been shown to establish long-range chromatin interactions (e.g.^67–69^). Whether PRC2 directly represses genes like *TPRX1* or acts via some other mechanism remains to be determined; iii) is the coordinate expression of LINE1 from its various loci across most/all chromosomes required for nuclear compartmentalization and 8C program repression? The repeated nature of LINE1 in the genome and the association of its RNA at chromatin with repressive factors, such as KAP1^7^, YTHDC1/SETDB1^70^ or the HUSH complex^71^ makes it an ideal genomic feature to coordinately silence gene expression programs. Alternatively, it is possible that specific LINE1 elements in unique chromosomal locations are critical for nucleolar organization and suppression of the 8C program. Moreover, it is worth keeping in mind that LINE1 RNA as well as some 8C loci such as *H3.XY* are also enriched at the nuclear lamina. In this regard, recent findings point to a remarkably dynamic nuclear lamina in mouse embryos at the 2C stage that may regulate chromatin organization at ZGA^72^. A systematic analysis of the unique landscape of nuclear compartmentalization of 8CLCs may enable the dissection of the role of activators (such as TPRX1) and repressors (such as LINE1) of the 8C state.

## Methods

### Primed hESC culture

H9 and H1 primed hESC lines^73^ were cultured in mTeSR Plus Medium (STEMCELL # 05825) on hESC-Qualified Matrigel-(VWR # CA89050-192) coated plates at 37 °C, 5% CO2, 20% O2 conditions^26^. Cells were passaged every 4-5 days by performing a PBS wash, adding then removing ReLeSR (STEMCELL # 05873) and incubating at 37 °C for 5 min. mTeSR medium was then added and the cell plate was tapped gently to release colonies. Without pipetting up and down, the cell suspension was dropped into new pre-Matrigel-coated wells. When single-cell seeding was required, the cell suspension was pipetted up and down, and cells were counted and seeded at the desired numbers with the addition of 20 µM ROCK inhibitor Y-27632 (Stemgent # S1049).

### RSeT hESC culture

RSeT hESCs were cultured in RSeT Medium (STEMCELL # 05975) at 37°C, 5% CO2, and 5% O2 conditions^24^. Primed hESCs were converted to RSeT conditions and passaged when colony sizes reached ∼100 µm, following the RSeT Medium manufacturer’s instructions. RSeT hESCs were used for experiments within ten passages of conversion.

For RSeT+DT conditions, RSeT hESCs were supplemented with 10 nM DZNep (Selleck # S7120) and 5nM TSA (Vetec # V900931) for 48 hours, with the medium changed at 24h. Similarly, for RSeT+eDT conditions, RSeT hESCs were supplemented with 50 nM DZNep and 20 nM TSA as described^3^ for 48 hours.

### MEF culture

Primary MEFs (passage 3) were obtained from the Lunenfeld-Tanenbaum Research Institute (LTRI) ESC Facility. About 2 x 10^5 MEFs were thawed on 0.1% gelatin-coated plates and cultured in MEF medium (DMEM, high glucose, GlutaMAX™ Supplement, pyruvate (Life Technologies # 10569), 15% FBS (BioTechne # S12450), 0.1 mM MEM nonessential amino acids (Thermo Fisher Scientific # 11140050) and 100 U/mL penicillin-streptomycin (Thermo Fisher Scientific # 15140122)). MEF feeders were used within 4-5 days of thawing.

### 4CL and e4CL hESC culture

4CL hESCs were converted from primed hESCs according to Mazid et al.^3^. Briefly, 1-1.5 x 10^5 primed hESCs were seeded as single cells per well of a 6-well plate pre-seeded with MEF feeder cells in mTeSR medium supplemented with 10 µM ROCK inhibitor Y-27632 at 37 °C in 20% O2. After 24 hours, medium was switched to 4CL [1:1 mix of Neurobasal medium (Gibco #21103049) and Advanced DMEM/F12 (Gibco #12634028) supplemented with N2 (Gibco # 17502048) and B27 (Gibco # 17504044) supplements, sodium pyruvate (Life Technologies # 11360070), non-essential amino acids (Life Technologies # 11140050L), GlutaMAX (Gibco # 35050061), penicillin-streptomycin (Life Technologies # 15140122), 10 nM DZNep (Selleck # S7120), 5 nM TSA (Vetec # V900931), 1 µM PD0325901 (Stemolecule # 04-0006), 5 µM IWR-1 (Sigma # I0161), 20 ng/ml human LIF (STEMCELL # 78055), 20 ng/ml ACTIVIN A (Peprotech # 120-14E), 50 µg/ml L-ascorbic acid (Sigma # A8960) and 0.2% (v/v) Geltrex (Life Technologies # A1413301)] and kept at 37°C in 5% O2. The medium was changed daily. When colony sizes reached ∼100 μm, cells were incubated in a 1:1 mix of TypleE:0.5 mM EDTA for 5 min treatment. Without tapping plates, the TypleE:EDTA mixture was replaced with 1 ml 4CL media. After tapping the plate and pipetting up and down, single cells were reseeded on Geltrex-(Life Technologies # A1413301) pre-coated or MEF-pre-seeded 6-well plates. 10 µM Y-27632 was added for 24 h the day after each passage. Cells were used for experiments within 3-5 passages.

To convert 4CL cells to e4CL cells, 2-3 x 10^5 per 6-well 4CL cells were seeded in 4CL medium, supplemented 24 h later with 50 nM DZNep and 20 nM TSA^3^ (e4CL). After 5 days of culturing in e4CL medium, cells were processed for analysis.

### dCas9-KRAB hESC line

Plasmids were built based on the sleeping-beauty transposon system^27^, as indicated in Figure S2D using enzymes from NEB. In a first step, the amaxaGFP sequence of pT2-CAG-amaxaGFP (a gift of the Izsvák lab) was removed using the AgeI and BglII sites and replaced with an artificial oligo sequence containing multiple enzyme digestion sites (FW-ccggtGGCGCGCCCGGGTGATCAa; RV-tGGCGCGCCCGGGTGATCAagatc). Next, AgeI and NotI sites were used to clone in the hUbC-dCas9-KRAB-T2a-Puro sequence from the pLV-hU6-sgRNA-hUbC-dCas9-KRAB-T2a-Puro plasmid^29^ (Addgene # 71236), generating pT2-hUbC-dCas9-KRAB-T2a-Puro such that the insert is now flanked by sleeping beauty inverted/direct repeats.

Primed H9 hESCs were used to generate the dCas9-KRAB transgenic line. For one well of a 6-well plate containing 5 x 10^6 single cells, 1.5 µg pT2-hUbC-dCas9-KRAB-T2a-Puro and 0.5 µg pCMV(CAT)T7-SB100X (a gift of the Izsvák lab, expressing an SB100X hyperactive transposase), were mixed with 4 µl Lipofectamine Stem (Life Technologies # STEM00008), in 200 µl Opti-MEM I (Thermo Fisher # 31985070). 24 hours later, the mTeSR medium was changed, and 0.3 µg/ml Puromycin (Life Technologies # A1113803) was added. For details on transfection, see below. Eight days after puromycin treatment, colonies were picked and transferred individually into wells of a 24-well plate. Colonies were expanded, confirmed to remain puromycin-resistant and validated for dCas9 protein expression by IF. Primed dCas9-KRAB transgenic hESCs thus generated were converted to RSeT conditions as above and used for CRISPRi as described below.

### ASO, siRNA and CRISPRi gRNA design

Locked nucleic acid (LNA)-gapmeR ASOs were designed targeting the L1HS full-length LINE1 L1RP (Genbank AF148856). Candidate target sites were first selected using CRISPR-RT (CRISPR RNA-Targeting), a Cas13a-based RNA target screening tool to pre-filter against off-target effects^74^. The candidate targets were then screened by BLASTN against the human genomic and transcript database, sites of low or no “Query cover” and “Percent Identity” scores were retained. Candidate target sites were further ranked for high conservation in full-length LINE1 elements using L1base (http://l1base.charite.de/l1base.php)^75^. Lastly, selected target sequences were adjusted to LNA-GapmeR design recommendations with assistance from QIAGEN customer support. QIAGEN Negative control A Antisense LNA GapmeR (LG00000002) and a sense oligo (SO) sequence of the interORF LINE1 ASO were used as controls (Table S1). Both naked and 6-FAM (Fluorescein)-labeled versions of the ASOs were synthesized and used (see below).

siRNAs targeting *TPRX1*^3^, *TP53*^76,77^ and *H3.XY*^35^ were selected from the literature (Table S1). The Dharmacon siGENOME Non-Targeting Control siRNAs (D-001210-03) were used as negative control (Table S1). All siRNAs were 6-FAM (Fluorescein) labeled and ordered from Horizon Discovery.

CRISPRi gRNAs (Table S1) targeting the 5’ UTR of L1RP with low off-target scores were selected by combining the output of CRISPOR (http://crispor.tefor.net/) and CRISPick (https://portals.broadinstitute.org/gppx/crispick/public)^78,79^. Individual sgRNA thus designed, as well as non-targeting control gRNAs^80^ were synthesized with overhangs (sense: 5′-ACCG, antisense: 5′-AAAC) subcloned into the BsaI sites of pGL3-U6-sgRNA-EGFP (Addgene # 107721)^81^.

### Oligos or plasmids transfection

Transfection of either ASOs or CRISPRi gRNA plasmids was conducted using Lipofectamine Stem, per manufacturer’s instructions. Typically, per 6-well with 2 ml of medium, 4 µl of Lipofectamine Stem and 25 nM ASOs or 1 µg/ml of plasmid were mixed in 200 µl of Opti-MEM Medium for one well of a 6-well plate containing single-cell hESCs at a density of 2.5-4 x 10^5 cells per well. The transfection mixture was added directly while the cells were in suspension. At the same time, 0.7 µM Trans-ISRIB (Sigma # 5095840001), 5 µM Emricasan (Sigma # SML2227) and 10 µM Y-27632 were added to improve transfection efficiency and cell survival^82^; these were removed during media change the next day. Cells were collected for downstream analyses 48 hours after transfection.

For siRNAs transfection, or co-transfection of ASOs and siRNAs, DharmaFECT 1 (Dharmacon # T-2001-02) was used. The siRNA transfection was performed following the manufacturer’s instructions, similar to above, including the additions of Trans-ISRIB, Emricasan, and Y-27632. 40 nM siRNAs were transfected, alone or together with 25 nM ASOs, per well of a 6-well plate.

### RNA polymerase I inhibition

RSeT hESCs were plated at 4 x 10^5 cells per well of a 6-well plate and switched to RSeT+DT medium the following day. After 24 hours in RSeT+DT conditions, cells were treated with 0.25 µM of BMH-21 (Sigma # 5099110001), while DMSO was added to control wells. Cells were fixed after 8 hours for immunofluorescence or collected after 24 hours for RNA isolation (see below).

### EZH2 inhibition

RSeT hESCs were plated at 8 x 10^4 cells per well of a 12-well plate. After 24 hours, the medium was changed to RSeT+DT, RSeT+DT+UNC2400, RSeT+DT+UNC1999 or RSeT+TSA+UNC1999. UNC1999 and UNC2400 were added at 2.5 µM final concentration. Cells were collected after 48 hours for RNA isolation.

### Self-renewal assays

Primed hESCs were plated on Matrigel-coated 12-well plates as single cells at a density of 8 x 10^4 per well in a mTeSR-based Cloning Medium, in which CloneR (STEMCELL # 05888) was used per manufacturer’s instructions to improve single cell survival and colony formation. Y-27632 was added for the first day. Three hours after cell seeding, the transfection mixture with 10 nM ASOs, 1 µl Lipofectamine Stem, and 50 µl Opti-MEM was added. Six days later, colonies were stained with Alkaline Phosphatase (AP) Red Substrate Kit (Vector Laboratories # SK-5100). AP+ colonies were manually counted under the microscope. Representative images showing colony morphologies were acquired with a Zeiss Axio z1 inverted microscope.

RSeT and RSeT+DT hESCs do not survive at the low densities used for standard colony formation assays (above). Thus, colony area was quantified by ImageJ analysis of microscope images 48 hours after ASO transfection (as above).

### RNA Extraction and qRT-PCR

In primed and RSeT hESCs, FAM+ ASO-transfected cells were isolated by fluorescence-activated cell sorting (FACS) on Sony MA900. The ASO transfection in RSeT+DT hESCs was highly efficient, with consistently >80% FAM+ cells (data not shown). Accordingly, in RSeT+DT conditions, cells were directly collected from the plates for RNA extraction, without sorting. In the case of co-transfection of ASOs (non-FAM labeled) with FAM-labeled siRNAs, FAM+ cells were isolated by FACS.

RNA was extracted using the RNeasy mini kit (QIAGEN # 74104) or Direct-zol RNA Microprep/Miniprep Kits (Zymo # R2061, # R2051) with on-column DNAse digestion, following the manufacturer’s instructions. cDNA was generated using the SuperScript IV VILO Master Mix (Life Technologies # 11756050) and used for qPCR using PowerUp SYBR Green Master Mix (Life Technologies # A25777) on a QuantStudio™ 5 Real-Time PCR System (Thermo Fisher). Data were normalized to housekeeping genes (UBB and/or GAPDH), and the results were further plotted and analyzed using GraphPad Prism. qPCR primers are listed in Table S1.

### RNA-sequencing and data analysis

30-200 ng of high-quality RNA (equal amounts of RNA per sample for each experiment) was used. 3-4 replicates were processed per sample, except where indicated. Sequencing libraries were prepared using the NEBNext Poly(A) mRNA Magnetic Isolation Module (NEB # E7490S) and NEBNext Ultra II Directional RNA Library Prep Kit (NEB # E7760L), following the instruction manuals. The quality of RNA and resulting libraries was assessed on a Fragment Analyzer (Agilent). Sequencing was performed on an Illumina NextSeq500 with 75bp single-end reads at the LTRI Sequencing Facility.

The RNA-seq reads underwent quality control and trimming using Trim Galore! v0.4.0, and were subsequently aligned to the human reference genome (GRCh19) with TopHat2 v2.0.13^83^. Gene or repetitive element counts (using an annotation file from UCSC RepeatMasker) were obtained using the FeatureCounts function from the Subread package (v1.5.0)^84^. TopHat and FeatureCounts settings used were as previously described^85^. The raw counts table was analyzed in R with edgeR (v3.40.2) for generating the log2-normalized counts table and comparing gene expression changes between the treated and control samples. Low-count genes - with normalized counts per million (CPM) ≤0.2 for RNA-seq in RSeT and RSeT+DT hESCs and CPM ≤0.5 for RNA-seq in primed hESCs - were filtered out. Differential gene expression was exported as topTable, which was further analyzed using tidyverse v1.3.2 and plotted using ggplot2 v3.4.1. MA plots and volcano plots were generated to display differential gene expression relative to controls: genes with an adj.P value < 0.05 and absolute log2FC > 1.5 were considered to be significantly up- or down-regulated. The log2-normalized counts were batch-corrected by the ComBat_seq function in sva v3.46.0^86^, and then used for generating multidimensional scaling (MDS) plots with limma v3.54.1, and heatmaps with gplots v3.1.3. Genes were clustered with the setting of hclust (as.dist(1-cor(t(df), method=“spearman”)), method=“complete”). Gene Set Enrichment Analysis (GSEA)^87^ was performed with the fgsea v1.24.0 package using pre-ranked t-values from the topTable. Cell or embryo signature gene sets used in the analysis above are listed in Table S2. Sets of genes upregulated or downregulated (adj.P value < 0.05 and log2FC > log2(1) or log2FC < −log2(1)) were intersected with gene lists of each chromosome in R with the dplyr function, and percentages were calculated relative to the total number of genes in each chromosome.

The gene sets of embryonic stages were generated with hierarchial clustering of the averaged Penalized Kernel Matrix Regression (avg.PKMR) values of genes in each embryonic stage from the scRNA-seq Yan et al. study^22^ using gplots v3.1.3 (see code in GitHub entry). Clusters of genes that define specific stages or lineages (Figure S5M) were used for further analysis (Table S2). 8cell_genes and Morulae_genes were further defined from the 8cell_Morulae_1687 gene cluster, by filtering avg.PKMR > 20 first, and then filtering as 8C genes if avg.PKRM (8cell / Morulae) >1.5 or as Morulae genes if avg.PKMR (Morulae / 8cell) >1.5. The other 8C-genes and Pre8C-gene sets were reported by Yu et al.^4^. The 8cell_genes_Stowers set are genes highlighted in Taubenschmid-Stowers et al.^2^.

To analyze the expression of LINE1 sub-families, 8C-TEs^88^ and selected protein-coding genes during embryonic stages, the raw counts table from bulk RNA-seq by Hendrickson et al.^21^ was analyzed with edgeR (v3.40.2) in R. The raw counts of all genes were used to calculate normalization factors, which were applied to calculate read-depth normalized counts for repetitive elements. The normalized counts table from this bulk RNA-Seq dataset^21^ and the scRNA-Seq datasets^22,23^ of early-stage human embryos were sub-selected for the repeats indicated (Figures 1A and S1A-S1E) or select protein-coding genes (Figures 5E and S14) and plotted as heatmaps with gplots v3.1.3. See in GitHub entry for details.

### Single-cell RNA-sequencing and data analysis

100,000 FAM+ (SO-L1 or ASO-L1 transfected) RSeT+DT hESCs were sorted into HBSS+2%FBS on ice. Cells were centrifuged at 500x g for 4 min, resuspended in 100 µl HBSS+2%FBS and immediately processed on a 10x Genomics Chromium platform using the Single Cell 3’ v3 RNA-seq kit, targeting a yield of ∼12,000 cells per sample. Final libraries were sequenced on an Illumina NovaSeq6000 S1 flow cell (Illumina) to a depth of ∼94,000 reads/cell for ASO-L1 and ∼50,000 reads/cell for SO-L1.

The raw sequencing data were aligned to the human reference genome (GRCh38) using Cell Ranger (v7.1.0) with default parameters. Cells with less than 5000 genes, more than 100000 unique molecular identifiers (UMIs), or higher than 15% mitochondrial UMIs were removed from the downstream analysis. All mitochondrial genes were filtered out from the gene expression matrices, which were then normalized using the Seurat v4 SCTransform function^89^. Cell clustering and visualization were performed by FindClusters and RunUMAP functions in the Seurat package^89^, with top 20 Principal components analysis (PCA) dimensions.

To project the hESC dataset onto the UMAP of human embryo data, the embryonic dataset was reanalyzed with Seurat at first. In brief, gene expression matrices were normalized using the NormalizeData function, and the top 3000 most variable genes were identified. The data was visualized using the RunUMAP with parameters: dims = 1:8, seed.use = 2023, n.neighbors = 30, n.epochs = 100, min.dist = 0.5. The SO-L1 and ASO-L1 datasets were also normalized using the NormalizeData function and filtered for the top 2000 most variable genes. The anchors between the embryo dataset and our hESC dataset were identified using FindTransferAnchors, with parameters: dims = 1:10, and k.anchor = 100. The projections were performed using MapQuery.

The set of genes up-regulated upon ASO-mediated L1 KD from bulk RNA-seq was assessed using Enricher (https://maayanlab.cloud/Enrichr/), which predicted high enrichment for certain gene modules, such as the p53 pathway or PRC2 targets. Select gene sets from this Enricher analysis were plotted onto the UMAP.

### LAD-seq

Three biological replicates per sample were prepared for LAD sequencing using pA-DamID, with an LMNB1 antibody (Abcam # ab16048), as described^41^. The Dam-treated Control and LMNB1+pA-Dam treated samples were subjected to DNA extraction using the ISOLATE II Genomic DNA Kit (Bioline # BIO-52066).

0.5 µg of genomic DNA was used for library preparation following the DamID-seq protocol^90^, with minor modifications: we omitted the DpnII digestion step for the pA-Dam reaction, as described in the pA-Dam Seq protocol^41^. Adaptors and primers used in this study are listed in Table S1. At the last step, after ten cycles of PCR amplification, the PCR products (size 1.2-1.5 kb) were purified using 1.6x volume AMPure XP beads (Beckman Coulter # A63880) and assessed by Fragment Analyzer (Agilent), then pooled for 100 bp single-end sequencing on an Illumina HiSeq2500 machine with 1%PhiX spike-in.

### NAD-seq

Three biological replicates per sample were prepared for NAD-seq following the crosslinked method of nucleoli isolation described by Vertii et al.^40^.In brief, after serial sonication with a Misonix XL-2000 ultrasonic cell disruptor (10s each burst at full power) and re-suspension in high then low magnesium buffers, cell nucleoli were released and monitored under the microscope. The nucleolar-fraction DNA (NF-DNA) and the control whole-cell DNA (WC-DNA) were extracted using NucleoBond AGX-20 columns (Takara # 740544) and NucleoBond® Buffer Set III (Takara # 740603), following the manufacturer’s instructions. To verify nucleoli enrichment, 4.6 ng of WC-DNA or NF-DNA per reaction was used for qPCR using primers listed in Table S1. NADs sequencing libraries were generated using the TruSeq DNA PCR-free Library Preparation kit (Illumina # 20015962); 350 bp fragments were sonicated and AMpure bead size-selected following the manual’s instructions. The libraries were subjected to 150 bp paired-end sequencing on an Illumina NovaSeq6000 instrument.

### LAD-seq and NAD-seq data analysis

For both the pA-DamID-LAD and NAD sequencing, raw sequencing data files were mapped to the human reference genome (GRCh38) using bwa, the output of which was filtered using samtools 1.10 (by “samtools view -b -F 4 -q 10”). LAD or NAD peaks were called from the bam file of each replicate using epic2^91^ with parameters bin_size=10000, gap=10 for the LADs and bin_size=10000, gap=5 for the NADs. LAD or NAD peaks called from at least two replicates out of the three samples sequenced were considered for further analysis.

Bam files from above analysis were uploaded to the UCSC genome browser to inspect LADs and NADs across whole chromosomes. The final LAD or NAD peaks bed files were intersected with bedtools^92^ 2.27.1 for their co-LAD/NAD coordinates, and calculated in R with the dplyr function for the coverage length ratio of LADs, NADs and co-LADs/NADs in each chromosome. To calculate the enrichment of LINE1 and other repetitive elements in LAD and NAD datasets, the final LAD or NAD peaks bed files were intersected with each indicated repetitive element’s bed file subsetted from RepeatMasker using the Intersection function in bedtools; this was also applied to a random DNA set of equal number and averaged size to LADs or NADs generated using the Random function in bedtools. The enrichment score was calculated as the ratio of the percentage of the length of LADs or NADs occupied by the repeat element divided by the corresponding percentage in the respective random DNA set.

### Cross-Linking Immunoprecipitation (CLIP) and qPCR

Plates of RSeT+DT hESCs were washed once with cold PBS, then were placed in cold PBS on ice with the lid open and UV-irradiated once in UV stratalinker 2400 with 400 mJ/cm^2^. Cells were harvested, and nuclear extraction was performed using a standard Abcam protocol (https://www.abcam.com/protocols/nuclear-extraction-protocol-nuclear-fractionation-protocol).

Prior to extraction, Nucleolin, LaminB1 or control Rabbit-IgG antibodies (See Table S3) were pre-bound to 50 µl Protein A Dynabeads (Life Technologies # 10002D) and incubated rotating for 3-6 hours at 4 °C. Beads were collected on a DynaMag (Thermo Fisher Scientific # 12321D) and re-suspended in High Salt CLIP buffer [1 M NaCl, 50 mM Tris pH 7.4, 1 mM EDTA, 0.5% Sodium deoxycholate, 1% NP40, protease and RNase inhibitors] containing 500 ng/mL tRNA (Invitrogen # AM7119) and 1 mg/ml RNase-free BSA (Sigma Aldrich # A9418) to block for 30 min, then collected and used immediately in RNA immunoprecipitation. Nuclear extracts (10 x 10^6 cells per IP) were pre-blocked for 30 min with 20 µl of Protein A Dynabeads at 4 °C, 30 min, then incubated with antibody-bound blocked beads overnight at 4 °C. Beads were washed twice with 1 ml High Salt Buffer and then twice with 1 ml Proteinase K Buffer [100 mM Tris-HCl pH 7.4, 50 mM NaCl, 10 mM EDTA, DnaseI (Life Technologies # 18068015) and RNase inhibitors (Life Technologies #10777019)]. Proteinase K Buffer washes were done for 15 min each on a 37 °C shaker at 800 rpm. After washes, beads were resuspended in 100uL of Proteinase K Buffer and RNA was eluted via addition of 50 µg Proteinase K (Life Technologies # 25530049) and 1 hour incubation at 55 °C. RNA was extracted from beads using Trizol and standard phenol-chloroform extraction. The aqueous phase containing the RNA was loaded onto RNeasy mini columns (QIAGEN) with 2x volume of 100% ethanol and RNA was purified according to kit’s manual, including an on-column DNase I treatment. Purified CLIP RNA was then used to generate cDNA for qRT-PCR. Primers are listed in Table S1.

### Immunofluorescence (IF)

Sterilized coverslips were inserted in wells of a 12-well plate and pre-Matrigel coated if needed. hESCs, whether transfected or not, were plated on the coverslips as described above. Two days post seeding (except where otherwise indicated), they were washed once with PBS and then fixed in 4% Paraformaldehyde (VWR # CAAAJ61899-AP) for 8 min, followed by two PBS washes. The plate was sealed with parafilm and stored in PBS at 4 °C for later use.

Fixed cells were permeabilized with 0.2% Triton X-100 in PBS for 2 min on ice. After two PBS-T (PBS + 0.1% Tween-20) washes, cells were incubated in blocking buffer [PBS-T + 1% BSA (Sigma Aldrich # A9418) + 10% Donkey serum (Sigma Aldrich # D9663)] for 30 minutes at room temperature (rt). Subsequently, cells were incubated with the primary antibodies diluted in the blocking buffer, washed 3 times in PBS-T for 5 min, followed by secondary antibody incubation (Table S3) and another set of PBS-T washes. Blocking and antibody incubations was performed on parafilm, with the coverslips facing down, and the wash steps were performed in the plates with the coverslips facing up. Coverslips were mounted onto slides with the VECTASHIELD® Antifade Mounting Medium with DAPI (Vector Laboratories # VECTH1200) and kept at rt for at least 1 hour before sealing with nail polish.

Most IF Images were acquired with an Andor BC43 Benchtop Confocal Microscope, except for the H3.XY and TPRX1 co-IF in e4CL-8CLCs (Figure S1F) and the Cas9 IF in dCAS9-KRAB hESCs (Figure S2D), which were acquired with on an Olympus Optigrid M Structured Illumination Microscope.

### RNA-FISH and RNA-FISH/IF

Cells were plated as above for the IF protocol, and fixed in 4% PFA for 15 minutes, followed by one PBS wash and two washes with ice-cold 70% Ethanol in DEPC-treated water. Coverslips were stored in 70% Ethanol in a parafilm-sealed plate at 4 °C for later use.

To perform RNA-FISH, cells were rehydrated in RNA wash buffer (WB) [10% Formamide + 2x SSC in DEPC water, prepared freshly] at rt for two 5 min incubations. Coverslips were then transferred to a piece of parafilm in a 15cm dish and incubated face down cell on RNA-FISH probe solution. Probes were designed and generated by Stellaris (see Table S4) to a stock concentration of 12.5 µM and were diluted 1:150 in hybridization buffer. To prepare the hybridization buffer, dextran sulfate (Sigma # D6001) was first dissolved in DEPC water at 10%, to which 1 mg/ml E.coli tRNA (Sigma # 10109541001), 2X SSC, 0.02% BSA, and 2 mM VRCs (NEB # S1402S) were added. After sterilizing with a 0.02 µM filter, the mixture was aliquoted into 475 µl per tube and frozen at −20 °C. The hybridization buffer was ready to use after adding 75 µl of formamide per aliquot freshly after thawing. Samples were incubated in the dark with probe solution, first for 45 min at 37 °C in a hybridization oven, then at 30 °C overnight without sealing the petri dish with parafilm. The next day, the dried coverslips were rehydrated by adding 50-100 µL of WB at the edges and waiting for a few minutes. Coverslips were transferred to a 12-well plate with WB for 5 min, followed by a 1-hour incubation in WB with 2.5 µl/ml RNAseOUT (Life Technologies # 10777019) at 37 °C. Coverslips were mounted onto microscope slides with VECTASHIELD® Antifade Mounting Medium with DAPI and then sealed with nail polish.

To perform RNA-FISH and IF co-staining, the RNA-FISH steps above were carried out with the following modifications. Primary antibodies were added with the probe in the hybridization buffer at 1:100 dilution and incubated in a humified container, in the dark first for 4 hours at 37 °C, then at 30 °C overnight. The next day, all steps being carried out in the dark, coverslips were transferred to a 12-well plate with WB for 5 min, then transferred to a new well containing WB with 2.5 µl/ml RNAseOUT and 1:200 diluted secondary antibodies, followed by a 1-hour incubation at 37 °C. After two 5 min washes in WB (with 0.5 µl/ml RNAseOUT) at rt, coverslips were mounted onto microscope slides with VECTASHIELD® Antifade Mounting Medium with DAPI and kept at rt for at least 1 hour before sealing with nail polish. RNA-FISH images were acquired on an Andor BC43 Benchtop Confocal Microscope, except for Figures 1C and S1H, which were acquired on a Leica DMI 6000 Spinning Disc Confocal Microscope.

### DNA-FISH/IF

DNA-FISH was performed essentially using the method described by Bolland et al.^93^, with modifications to perform DNA-FISH and IF co-staining by combining it with the method of Chaumeil et al.^94^.

DNA-FISH probes were generated from BACs (listed in Table S4) or LINE1 DNA plasmids. To enrich for signal from full-length LINE1 elements, a hL1-5’_3.3kb plasmid was generated by sub-cloning the 5’ end ∼3.3kb LINE1 sequence from the EF06R plasmid (Addgene # 42940)^95^ to the backbone of the pU6-sgRosa26-1Cbh-Cas9-T2A-BFP plasmid (Addgene # 64216)^96^, using the NotI and XbaI sites. BAC DNA or hL1-5’_3.3kb plasmids were purified using the NucleoBond® Xtra Midi Plus EF kit (Takara # 740422.50) following instructions for low-copy or high-copy plasmids, respectively. 10 µg BAD or hL1-5’_3.3kb plasmids were nick-translated with DNase I (Sigma # 04716728001) and DNA Polymerase I (NEB # M0209S). 150bp-700bp DNA fragments were generated and incorporated with Aminoallyl-dUTP (Invitrogen™ # AM8439), then purified and subjected to fluorescent dye coupling as described^93^ (see Table S4).

Single cells were collected as above (ASO-transfected cells were collected for FAM positivity by FACS), and plated in their respective medium on Poly-Prep Slides (Sigma # P0425-72EA) in an area encircled with a hydrophobic pen and pre-coated with Matrigel. After three hours in the incubator, the cells were fixed, permeabilized, washed and stored in 50% glycerol in PBS at - 20 °C, as described^93^.

For DNA-FISH/IF, slides were thawed in 20% glycerol/PBS at rt and subjected to 3 cycles of freeze/thawing in liquid nitrogen, followed by two PBS washes and permeabilization in 0.5% saponin/0.5% Triton X-100/PBS for 30 min at rt. Next, 100 µl blocking solution (2.5% BSA/10% Donkey serum/PBS-T) was added to the cell area and covered with a coverslip, followed by a 30 min incubation. The coverslip was then gently removed, and immediately 100 µl of a 1:100 dilution of primary antibody in blocking buffer was added, followed by an incubation at rt for 2 hours. After 3 washes in 0.2% BSA/PBS-T solution, cells were incubated in 100 µl of a 1:200 dilution of secondary antibody in the dark for 1 hour (see Table S3 for antibodies used). After the IF steps, cells were washed 3 times in PBS-T and fixed in 2% PFA for 10 minutes at rt, followed by a 30 min incubation in 0.1 M HCl, a PBS wash, and 30 min permeabilization in 0.5% saponin, 0.5% Triton X-100 in PBS. After two PBS washes and equilibration in 50% formamide in 2x SSC, cells were ready for addition of DNA-FISH probes mixture.

The DNA-FISH probe mixtures were prepared as described^93^ during the cell permeabilization and IF steps. The Chromosome 19 probe (Creative Bioarray # FWCP-19) mixture was treated with 7 min 80°C denaturation treatment followed by a 10 min 37°C incubation and then directly added to the slides, per manufacturer’s instructions.

When cells and probes were both ready, the slides were briefly drained with Q-tips and immediately covered with 10 µl of probe mixture that were pre-dropped on a coverslip. After sealing the coverslips with rubber cement (Marabu Fixogum), slides were heated at 78 °C for 2 min on a hot plate. Slides were then incubated overnight at 37 °C in a dark and humidified chamber. The next day, the slides were subjected to a series of wash steps following the protocol by Bolland et al.^93^., then mounted with VECTASHIELD® Antifade Mounting Medium with DAPI and kept at room temperature for at least 1 hour before sealing with nail polish. The DNA-FISH and IF co-staining images were acquired on an Andor BC43 Benchtop Confocal Microscope.

### Image quantification

ImageJ was used for quantifying signal intensity, area sizes and distances, as detailed in figure legends. The area of the nucleus or the nucleolus was manually traced based on DAPI or NCL/B23 signal. Quantification of the number of LINE1 foci per cell was acquired at threshold of 5 using a published RNA-FISH quantification method^97^.

The DNA-FISH/IF images were analyzed using Imaris. Chr.19, NCL (nucleolus), DAPI (nucleus), and TPRX1 DNA loci were 3D reconstructed using the Create Surface function in Imaris to identify their territories and location, and quantify their overlapping volumes. Representative cells and their 3D reconstructed territories were exported in supplementary videos.

### Statistical analyses

The image quantification data are presented as mean ± SEM and analyzed with GraphPad Prism v9.3.1 or R v4.0.3., using student t-test. Depending on the data distribution or comparisons being made, the t-test methods are selected differently, and noted in each figure legend.

RT-qPCR and CLIP-qPCR are plotted for the averaged fold changes ±SEM of n times biological replicates with GraphPad Prism. Ratio-paired student’s t-test was used for statistical analysis.

For RNA-seq data analysis, differential gene expression was calculated as described above. Comparison of expression changes between genes sets were displayed by violin plot and analyzed for statistical significance using the Wilcoxon test in R.

Sample size, number of replicates of studies, and statistical tests were chosen based on experience and variability of in vitro studies and stated in each figure legend. Unless otherwise indicated, all experiments were repeated at least three independent times.

### Data and code availability

Sequencing data, including raw reads and processed data (raw counts, normalized counts and differential gene expression table) have been deposited on the NCBI Gene Expression Omnibus repository (GEO, http://ncbi.nlm.nih.gov/geo) and will be accessible upon publication. The data reference Series for this study is GSE232939.

scRNA-Seq from this study was compared to the scRNAseq data from human embryos^30^ and e4CL-8CLCs^3^, which are available under the accession numbers E-MTAB-3929 and CNP0001454, respectively.

Raw-counts tables or normalized-counts tables from other studies of human embryos using scRNA-seq or bulk RNA-seq studies^21–23^ were also used in this work. Processed data were downloaded from the supplementary tables in these publications.

The authors declare that all other data supporting the findings of this study are available within the paper and its supplementary information files. Code supporting this study is available at a dedicated Github repository: https://github.com/tzzhangjuan1/LINE1_hESC.

## Acknowledgments

We thank members of the Santos lab, M. Lupien and M. Percharde for critical reading of the manuscript. We thank all members of Santos lab for their input throughout the project, in particular E. Collignon and T. Macrae for guidance on bioinformatics analyses. We are grateful to the J. Ellis and B. Di Stefano for assistance with hESC cell culture, to Z. Izsvák and J. Wang for sharing plasmids, to C. Arrowsmith and the Structural Genomics Consortium for sharing PRC2 inhibitors and to P. Maass, M. Lupien, L. Pelletier, B. van Steensel, T. van Schaik, M. Percharde and Y. Yin for precious advice and technical assistance. We are also thankful to K. Chan at the LTRI Sequencing Core, A. Bang at the LTRI Flow Cytometry Facility, R. Bielecki, L. Brown at the LTRI Microscopy Facility and M. Kownacka at the LTRI ESC Facility for their core facility support. This work was supported by a 2022 Medicine by Design Research Application Support Initiative (RASI) Award to S.S., the Natural Sciences and Engineering Research Council of Canada (RGPIN-2022-05134) to M.M.H., and a Canada 150 Research Chair in Developmental Epigenetics, the Great Gulf Homes Charitable Foundation, and a 2019 Medicine by Design New Ideas Award to M.R.-S..

## Author contributions

M.R.-S. and J.Z. conceived of the project and designed the experiments. J.Z. performed majority of the experiments and interpreted the data. L.M. performed CLIP, and the CRISPRi experiments with a cell line generated by J.Z.. K.M. conducted Pol I inhibition experiments. D.P.C., D.T., and J.L.W. provided guidance with scRNA-seq and prepared the libraries. L.W. analyzed and interpreted the scRNA-seq data, with assistance from M.A.M. and M.A.E.. M.A.E. provided plasmids and relevant advice regarding 4CL and e4CL cultures. L.H. and M.M.H. processed the LADs-seq and NADs-seq data, and L.H and J.Z. performed downstream analysis. S.S. and K.T. assisted J.Z. in IF and DNA-FISH co-staining experiments. M.R.-S. supervised the project. J.Z. and M.R.-S. wrote the manuscript with input from all authors.

## Supplemental information

**Table S1. Sequences of ASOs-, siRNAs-, CRISPRi-gRNAs, primers and adaptor oligos used in this study.**

**Table S2. Gene sets used in this study**

Notes: The gene sets from scRNA-seq^22^ are re-derived here by hierarchical clustering. See STAR Methods and Figure S5M for details.

**Table S3. Antibodies used for Immunofluorescence (IF) and Cross-Linking Immunoprecipitation (CLIP)**

**Table S4. Sequences of RNA-FISH probes and list of plasmids for DNA-FISH probes**

**Videos S1-S2.** Representative videos of chromosome 19 by DNA-FISH and co-IF staining with NCL antibody in e4CL-induced 8CLCs and 4CL-cultured naïve hESCs. Corresponds to representative cells in Figure 5D. The Chr.19 and NCL-marked nucleolus territories imaged by microscope are three-dimensionally reconstructed in Imaris using the Create surface function. Scale bar, 20 µm.

**Video S3-S6.** Representative videos of *TPRX1* loci by DNA-FISH and co-IF staining with B23, and H3.XY (e4CL cells) or LMNB1 (4CL, RSeT and Primed hESCs). Corresponds to representative cells in Figure S10B, indicated by large yellow arrows. The B23-marked nucleolus territories and DAPI-marked nucleus regions are three-dimensionally reconstructed in Imaris using the Create surface function. Scale bar, 20 µm.

---

**Figure S1.**
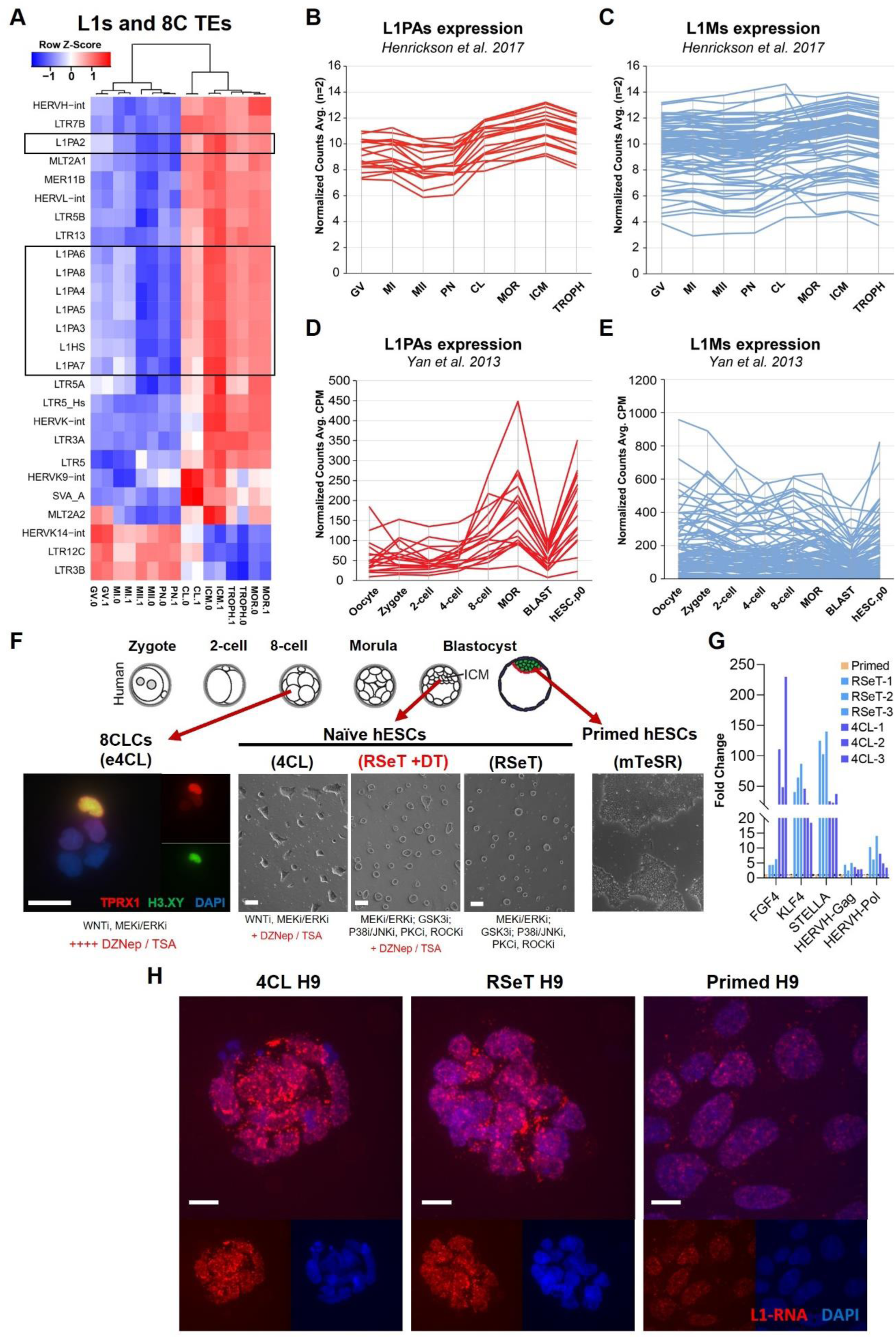
LINE1 RNA is highly expressed in early embryos and hESCs. (A) Heatmap showing induction of primate-specific LINE1 subfamilies and 8C-morula expressed TEs^88^ in RNA-seq datasets from developing human embryos^21^. (B and C) Average normalized counts of n=2 independent RNA-seq samples^21^ showing induction of primate-specific L1PA subfamilies (B), but not of older L1M subfamilies (C) during human embryonic cleavage stages. GV, germinal vesicle; MI, metaphase I; MII, metaphase II; PN, pronuclear stage; CL, cleavage; MOR, morula; ICM, inner cell mass; TROPH, trophectoderm. (D and E) Average normalized counts from scRNA-seq^22^ data showing induction of L1PAs (D) but not L1Ms (E) during early human embryonic development. MOR, morula; BLAST, blastocyst. (F) Schematic of hESCs culture conditions used in this study, and their corresponding embryo stages in vivo. DZNep and TSA, key components in the 4CL media^3^, were supplemented to the RSeT medium for 48 hours. IF staining of TPRX1 and H3.XY identifies 8CLCs induced by e4CL medium^3^. IF images, scale bar, 20 µm. Cell colony images, Scale bar, 100 µm. (G) qRT-PCR showing strong induction of markers of the naïve state in both RSeT and 4CL hESCs (3 batches each), relative to Primed hESCs. (H) RNA-FISH in 4CL, RSeT and Primed hESCs showing predominant nuclear localization of LINE1 RNA in hESCs. Representative of at least two independent experiments. Scale bar, 10 µm.

**Figure S2.**
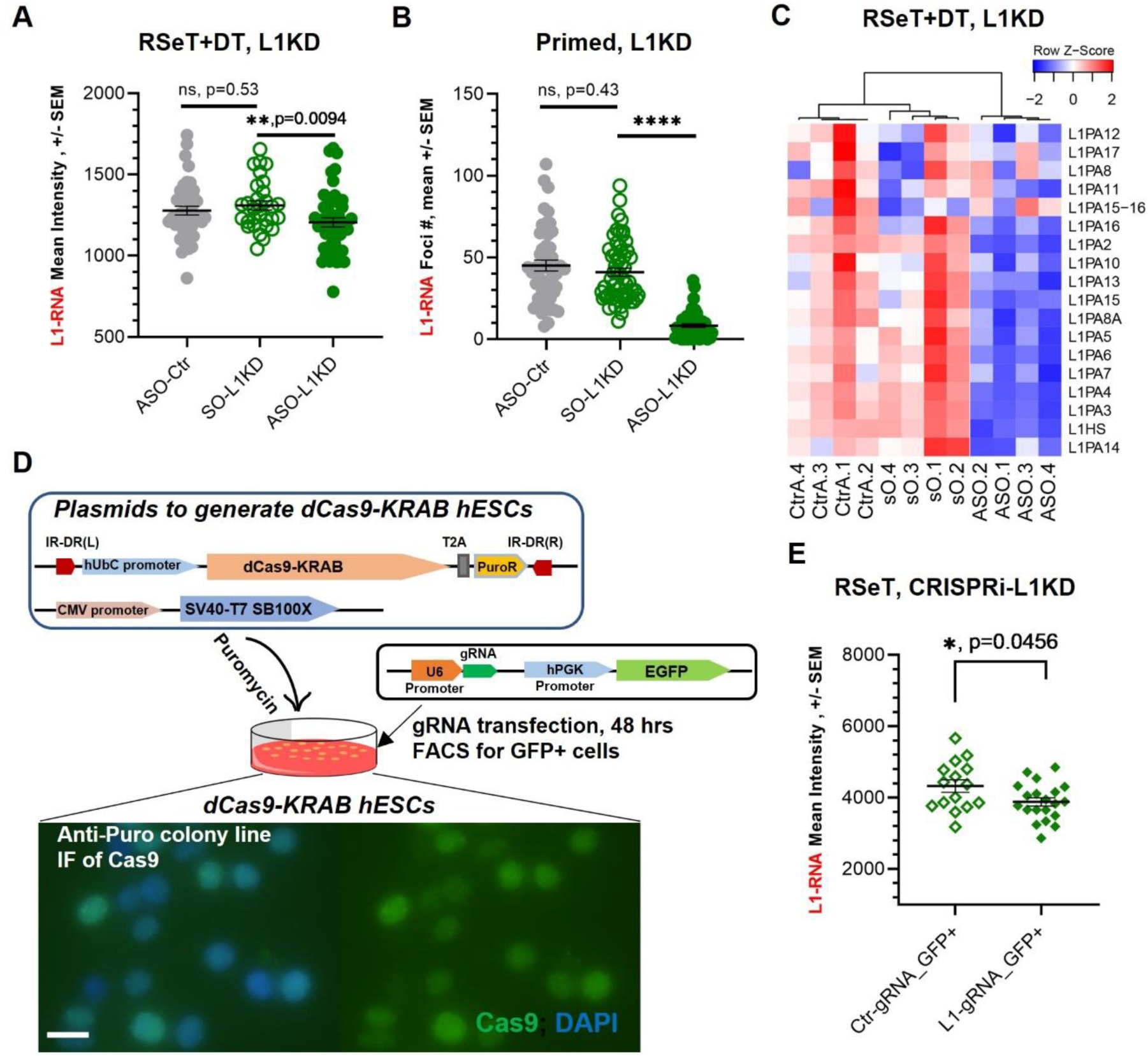
ASOs and CRISPRi efficiently knockdown L1 RNA. (A) Quantification of LINE1 RNA-FISH signal intensity in RSeT+DT hESCs after 48 hours of ASO-L1, SO-L1, or ASO-Ctr transfection, showing efficient knockdown in ASO-L1 transfected cells compared to controls. Representative of two independent experiments. Welch’s t-test. ns = p > 0.05, ** = p < 0.01. (B) Quantification of LINE1 RNA-FISH foci per cell in primed hESCs after 48 hours of ASO-L1, SO-L1, or ASO-Ctr transfection, indicating significant reduction of LINE1 in ASO-L1 transfected cells compared to controls. Representative of at least four independent experiments. Mann-Whitney test. ns = p > 0.05, **** = p < 0.0001. (C) Heatmap showing reduction of L1HS and L1PAs upon ASO-L1 KD in hESCs compared to SO-L1 and ASO-Ctr controls. (D) Schematic of generation of the CRISPRi system in hESCs using the Sleeping Beauty transposon system (see Methods for details). After puromycin selection and expansion of resistant colonies, Cas9 expression was validated by IF staining (bottom panel). Scale bar, 20 µm. (E) Quantification of LINE1 RNA-FISH signal intensity in dCas9-KRAB hESCs 48 hours after transfection of L1-gRNA or Ctr-gRNA plasmids. The LINE1 RNA intensity in the L1-gRNA transfected EGFP+ cells is significantly reduced compared to control. Representative of two independent experiments. Welch’s t-test. * = p < 0.05.

**Figure S3.**
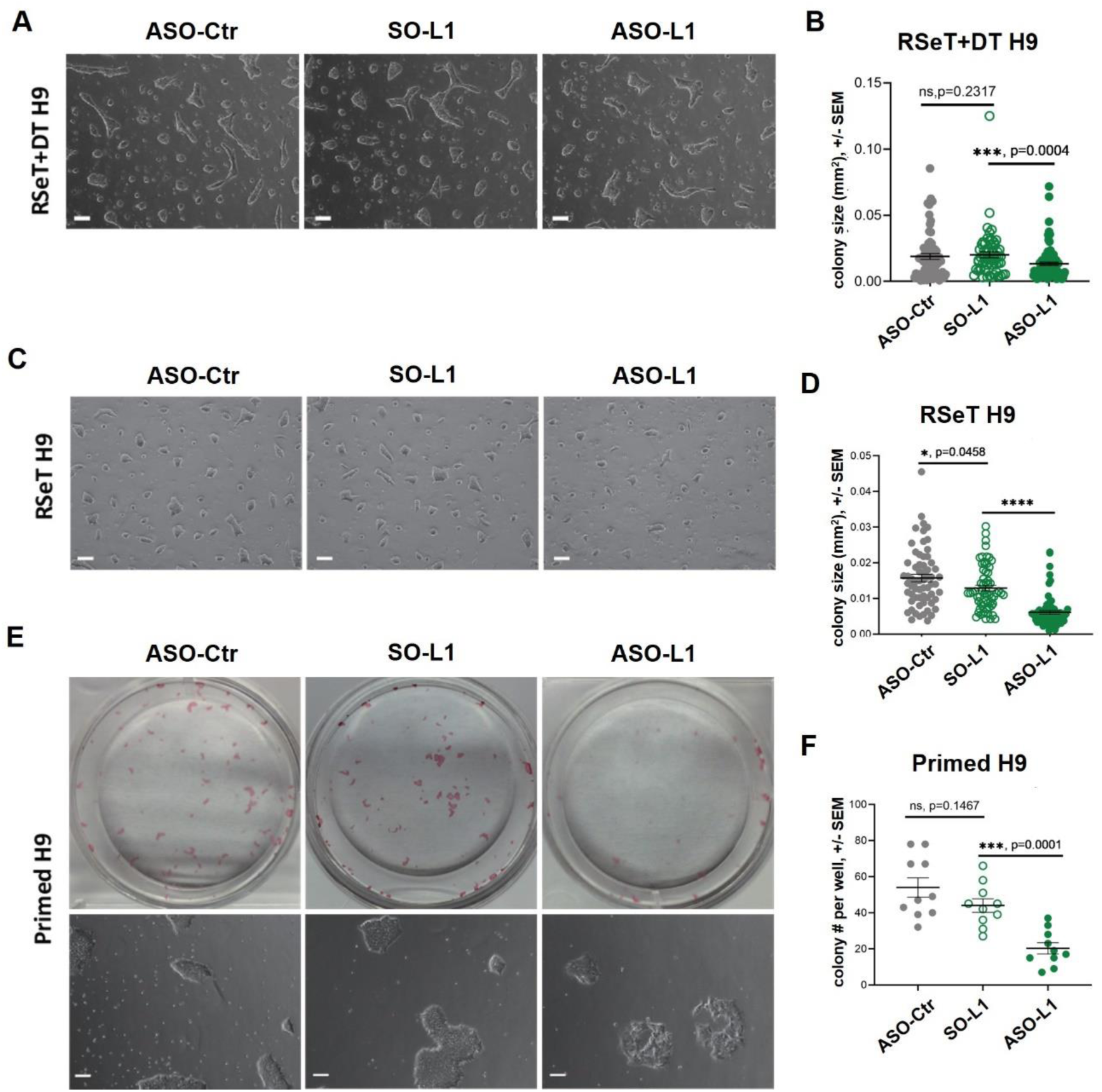
LINE1 KD impairs self-renewal of hESCs. (A-D) Colony size analysis in (A and B) RSeT+DT and (C and D) RSeT hESCs, respectively. ASO-L1 transfected cells display significantly smaller colony sizes than control ASO-Ctr and SO-L1 transfected cells. Representative images on the left (A and C), quantified on the right (B and D). Representative of three independent experiments. Mann-Whitney test. ns = p > 0.05, * = p < 0.05, *** = p< 0.001, **** = p < 0.0001. Scale bar, 100 µm. (E and F) Colony formation assay in primed hESCs. ASO-L1 transfected cells generate significantly less colonies than control ASO-Ctr and SO-L1 transfected cells. AP-stained (upper panel in E) colony numbers per 12-well were quantified and plotted in (F). ASO-L1 transfected primed hESCs also display abnormal colony morphology (lower panel in E). Representative of three independent experiments. Mann Welch’s t-test. ns = p > 0.05, *** = p< 0.001. Scale bar, 100 µm.

**Figure S4.**
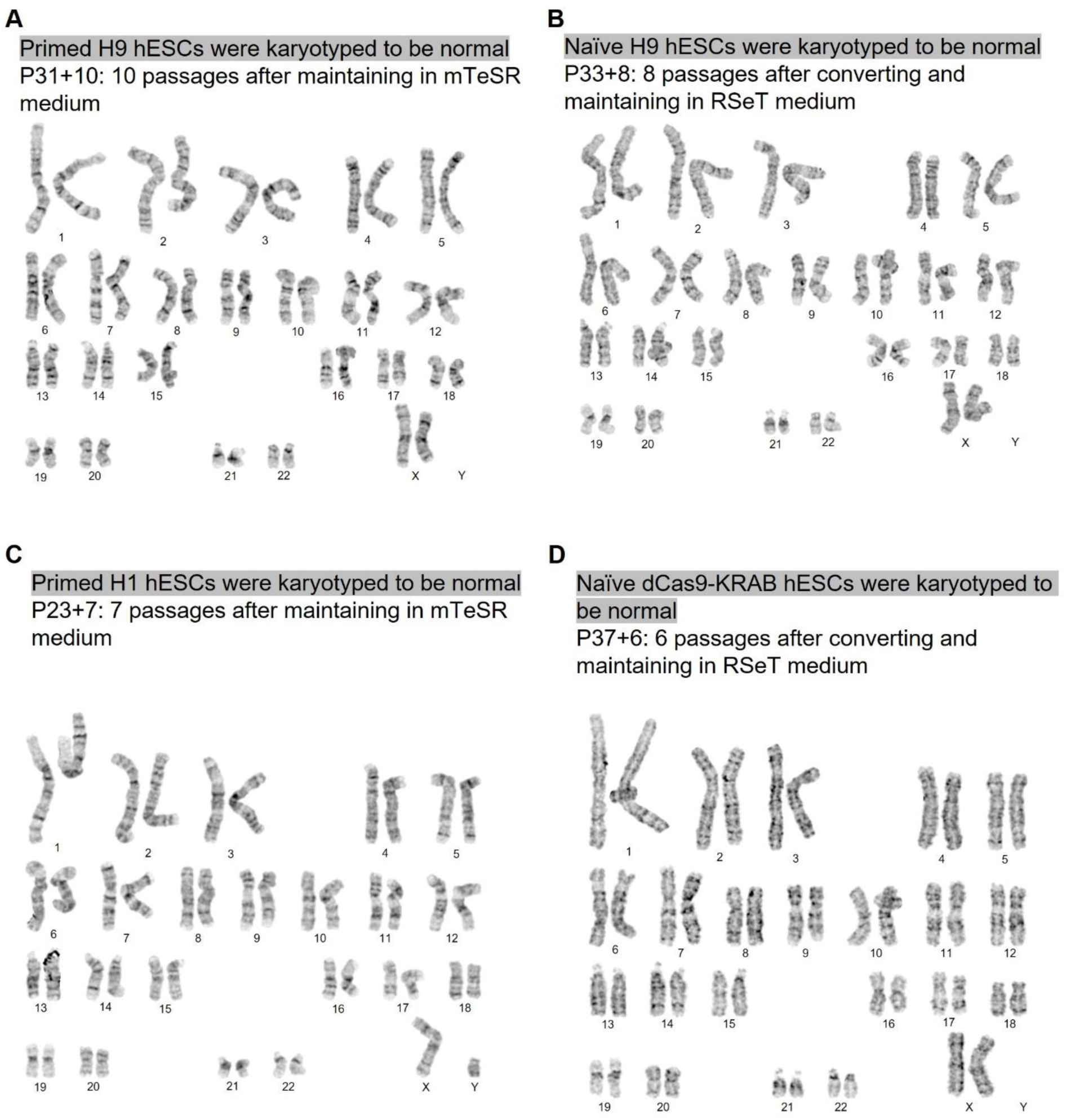
hESC lines used in this study display a normal karyotype. (A and B) Karyotyping of H9 hESCs maintained in (A) mTeSR primed and (B) RSeT naïve-like conditions. Cells are confirmed to have a normal karyotype. (C) Karyotyping of primed H1 hESCs. Cells are confirmed have a normal karyotype. (D) Karyotyping of the dCas9-KRAB transgenic H9 hESC line maintained in RSeT naïve-like conditions. Cells after dCas9-KRAB sequence insertion is confirmed to be normal karyotype.

**Figure S5.**
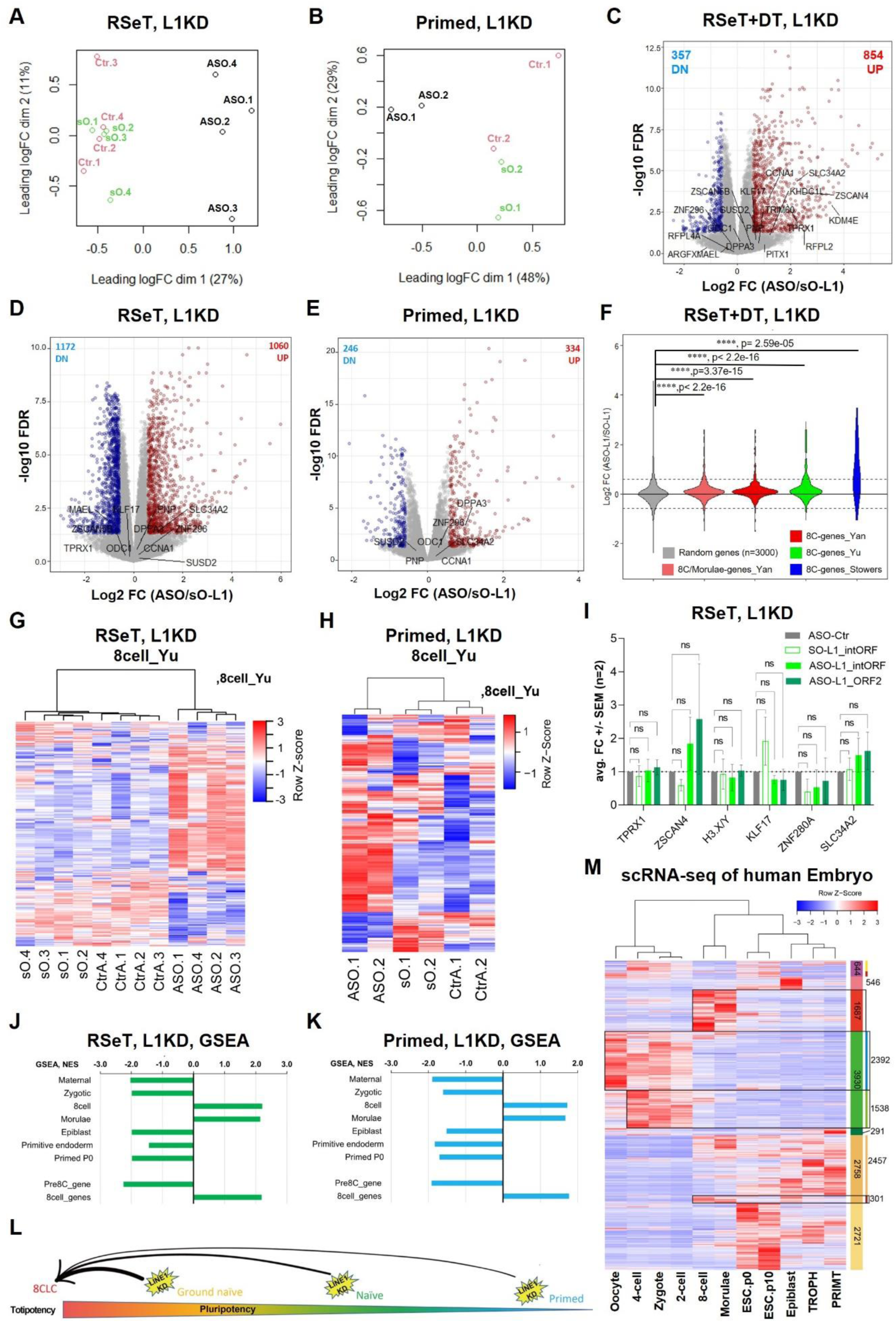
LINE1 KD disrupts the transcriptional profile of hESCs and induces 8C signatures. (A and B) MDS plot of genes across all samples, showing that ASO-L1 KD (A) RSeT and (B) primed hESCs have distinct gene expression profiles from their respective control SO-L1 and ASO-Ctr samples. (C-E) Volcano plot showing log2-fold change (FC) in gene expression following ASO-L1 KD in (C) RSeT+DT, (D) RSeT and (E) Primed hESCs compared to control SO-L1 transfected cells. Red or blue highlight genes of adj.P-value < 0.05 and FC > 1.5 or < −1.5, respectively. 8C marker genes from Taubenschmid-Stowers et al.^2^ are labeled. (F) Violin plot of data from ASO-L1KD in RSeT+DT hESCs showing significant upregulation of morulae and 8C gene sets from different studies^22,2,4^, compared to a random set of n=3000 genes. Wilcoxon test. **** = p < 0.0001. (G and H) Heatmap showing induction of gene sets of the 8C stage from X. Yu et al.^4^ in (G) RSeT and (H) Primed hESCs upon ASO-L1KD compared to ASO-Ctr and SO-L1 controls. (I) qRT-PCR showing a lack of upregulation of key 8C marker genes in LINE1 KD RSeT H9 hESCs (without DT) with ASOs targeting the inter-ORF and ORF2 sites, respectively. Data are mean ± SEM, n = 2 biological replicates. Ratio paired Student’s t-tests. ns = p > 0.05, (J and K) Gene Set Enrichment Analysis of the transcriptional profile of ASO-L1 KD (J) RSeT and (K) Primed hESCs for the enrichment of gene sets from different stages of pre-implantation development and 8C/pre-8C gene sets^4,22^. (L) Schematic illustrating the impact of LINE1 KD in hESCs of distinct naïve states. hESCs of a higher naïve nature (RSeT+DT > RSeT >Primed) are more permissive for derepression of the 8C program upon KD of LINE1. See Figures 1F and S5J-S5K. (M) Heatmap analysis of scRNA-seq data from early human embryos^22^, see Methods for details. Gene sets of each human embryonic stage are hierarchy clustered and listed in Table S2. TROPH, trophectoderm; PRIMT, Primitive endoderm.

**Figure S6.**
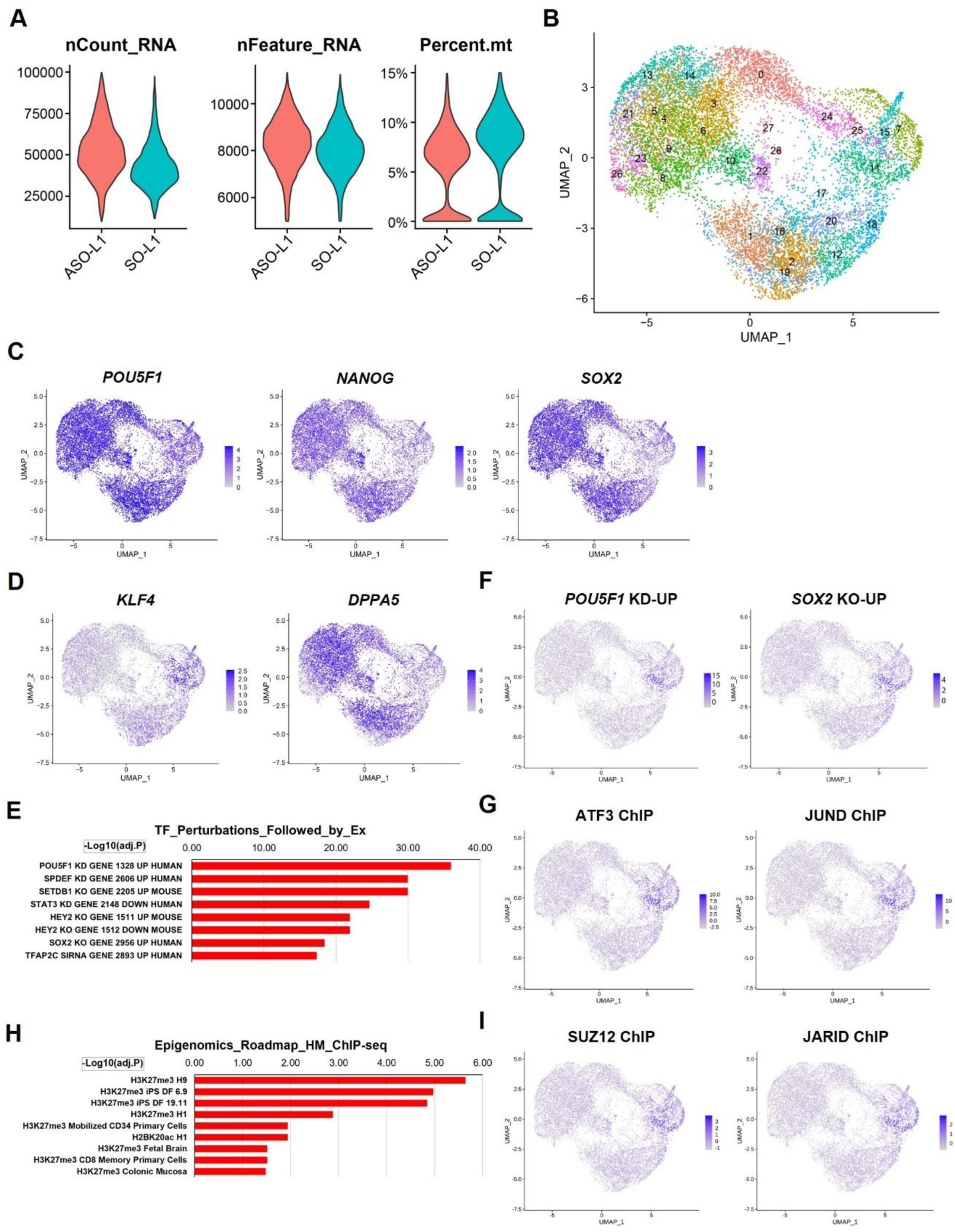
Additional analyses of scRNA-seq of LINE1 KD hESCs. (A) Metrics assessing the quality of scRNA-seq libraries, plotting transcripts per cell (nCount_RNA), genes detected per cell (nFeature_RNA) and mitochondria reads percentage (percet.mt) in the ASO-L1 KD and SO-L1 samples. (B) Visualization of 28 cell clusters with uniform manifold projection (UMAP). Cluster 15 is identified to as 8CLCs cluster. (C and D) UMAP view of cellular expression levels of (**c**) pluripotency markers; (**d**) naïve hESC markers. (E) Top signatures of genetic perturbations enriched at genes upregulated upon L1KD in RSeT+DT hESCs (see Methods). (F) UMAP view of expression levels of genes induced by *POU5F1* KD or *SOX2* KO. (G) Top chromatin-bound factors enriched at genes upregulated upon L1KD in RSeT+DT hESCs (complementary to analysis shown in Figure 2E). (H) UMAP view of expression levels of stress-related gene targets of ATF3 or JUND. (I) UMAP view of expression levels of targets of PRC2.

**Figure S7.**
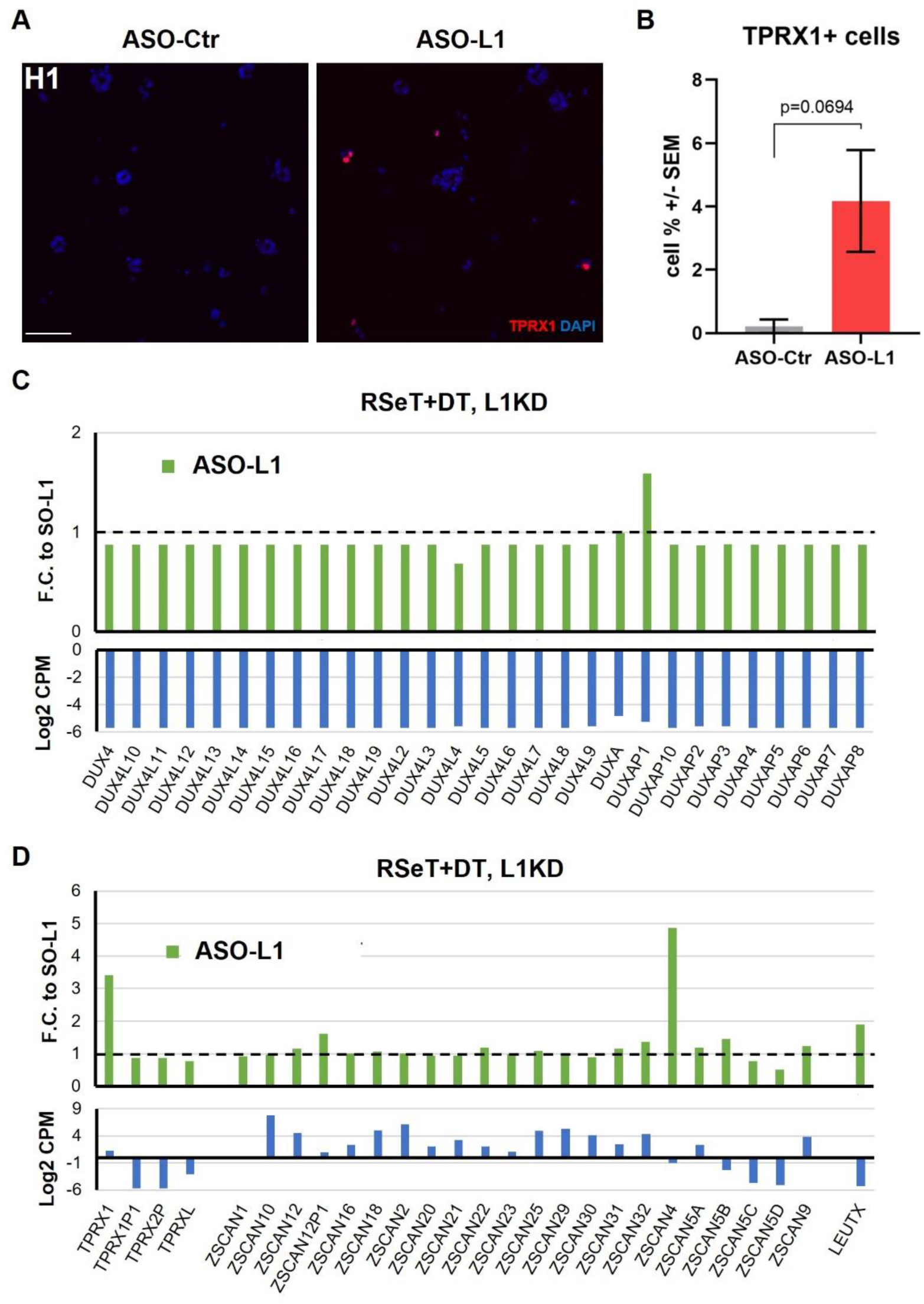
The expression of *TPRX1*, *DUX4* and *DUX4*-like genes in LINE1 KD hESCs. (A) IF of TPRX1 in ASO-L1 transfected H1 hESCs compared to ASO-Ctr control. Representative of two independent experiments. Scale, 100 µm. (B) Quantification of TPRX1+ cells ratio in (A). Same as in Figure 3B, 380∼400 cells per sample from five random views were quantified. Data are mean ± SEM. Welch’s t-test. Representative of two independent experiments. (C and D) Plot of expression of select genes from RNA-seq data of ASO-L1 KD RSeT+DT ESCs compared to SO-L1 control for (upper panel) fold change (F.C.) and (lower panel) normalized log2 counts per million (log2 CPM). These data indicate (C) undetectable expression of *DUX4* and *DUX4*-like genes, in contrast to (D) *TPRXs*, *ZSCANs*, and *LEUTX* genes.

**Figure S8.**
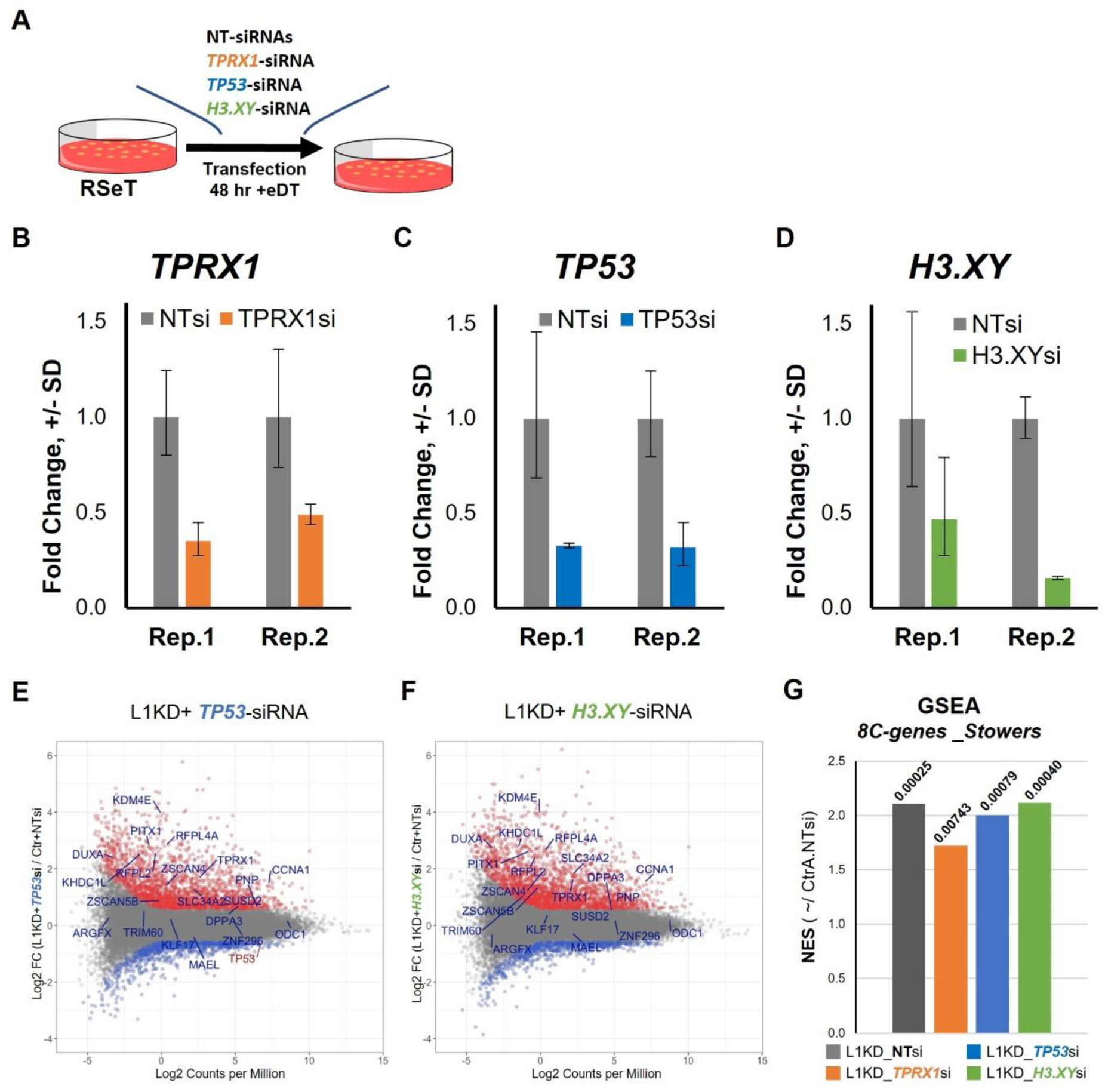
*TPRX1* KD partially rescues LINE1 KD-mediated 8C gene upregulation, whereas *TP53* and *H3.XY* KD has no detectable impact. (A) Schematic of siRNAs knockdown of *TPRX1*, *TP53*, and *H3.XY* in RSeT+eDT hESCs, compared with a control non-targeting (NT) siRNAs. RSeT+eDT conditions were used to induce a higher level of putative 8C regulators and better assess their siRNA-mediated KD (see STAR Methods). (B-D) qRT-PCR data of two biological replicates (Rep.) of siRNA KD in RSeT+eDT hESCs, as shown in **a**, validating efficient KD of corresponding targets: (B) *TPRX1*-siRNA (*TPRX1*si); (C) *H3.XY*-siRNA (*H3.XY*si); and (D) *TP53*-siRNA (*TP53*si). Data are mean fold changes of n=3 technical repeats, ± SD. (E and F) MA plot showing log2 fold changes in gene expression following (E) L1KD+*TP53*-siRNA and (F) L1KD+*H3.XY*-siRNA compared to control transfected cells (ASO-Ctr+NT-siRNA). Red or blue highlight genes of adj.P-value < 0.05 and FC > 1.5 or < −1.5, respectively. 8C marker genes from Taubenschmid-Stowers et al.^2^ are labeled in dark blue. (G) Plot of the NES value from the GSEA analysis showing reduced enrichment of 8C marker genes^2^ in L1 KD+*TPRX1*-siRNA transfected hESCs compared to L1 KD+NTsi-RNA transfected cells, while L1 KD+*TP53*-siRNA and L1 KD+*H3.XY*-siRNA transfected hESCs shows similar induction level of 8C genes. adj.P values of each sample by GESA analysis are indicated on top of the bar.

**Figure S9.**
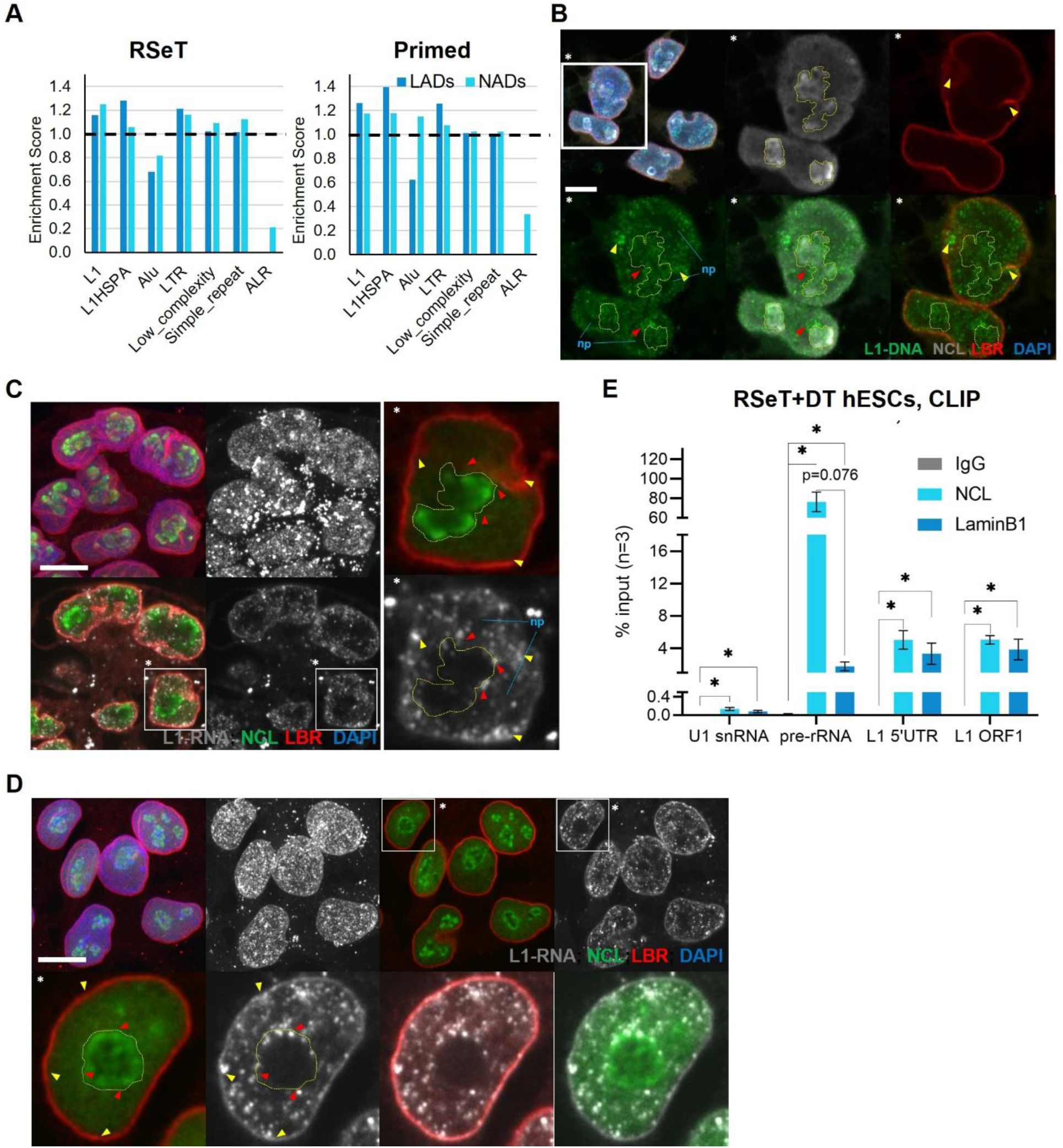
LINE1 elements are enriched at the nuclear lamina and the periphery of the nucleolus. (A) Enrichment of transposon families in LADs and NADs of RSeT (left panel) and Primed hESCs (right panel). Coverage ratio of each family in LADs or NADs was compared to the coverage ratio in a random set of genomic sequences of equal number and averaged size (set as 1). L1HSPA: L1HS and L1PA1-8 of length > 5.5 kb. (B) Representative image of LINE1 DNA-FISH and co-IF staining for NCL and LBR antibody in RSeT+DT hESCs, showing that LINE1 loci are enriched in the vicinity of nucleolar and laminar domains. Boxed area (**\***) in the max projected image (top left panel) is enlarged and displayed as a representative single z-stack image in separate channels. Yellow dotted line encircles the nucleolus area. Yellow and red arrows point to LINE1 loci-enriched spots at the nucleolar and laminar domains, respectively. Noted the nucleoplasm (np) areas are sparse in LINE1 loci. Representative of three independent experiments. Scale bar, 10 µm. (C and D) Representative image of LINE1 RNA-FISH and co-IF staining for NCL and LBR antibody in (C) RSeT+DT hESCs and (D) Primed hESCs, showing that LINE1 RNA are enriched in the vicinity of nucleolar and laminar domains. The max projected image (top left panels in C and two top left panels in D) displays high expression of LINE1 in the nucleus. A representative plane is shown at the bottom left panels in (C) and two top right panels in (D), with the boxed area (*) enlarged at the right panels in (C) and bottom panels in (D), with color channels separated. Yellow dotted line encircles the nucleolus area. Yellow and red arrows point to LINE1 RNA foci-enriched spots at the nucleolar and laminar domains, respectively. Noted the nucleoplasm (np) areas are sparse in LINE1 foci. Representative of at least three independent experiments. Scale bar, 10 µm. (E) Cross-linking immunoprecipitation (CLIP) qRT-PCR analysis for indicated RNAs pulled-down with NCL or LMNB1 antibodies, compared to IgG pulldown. LINE1 RNA is associated with both NCL and LMNB1. Note the high enrichment for pre-rRNA is highly in the NCL pulldown. Data are mean ± SEM, n = 3 biological replicates. Multi Ratio paired t-test. * = p < 0.05.

**Figure S10.**
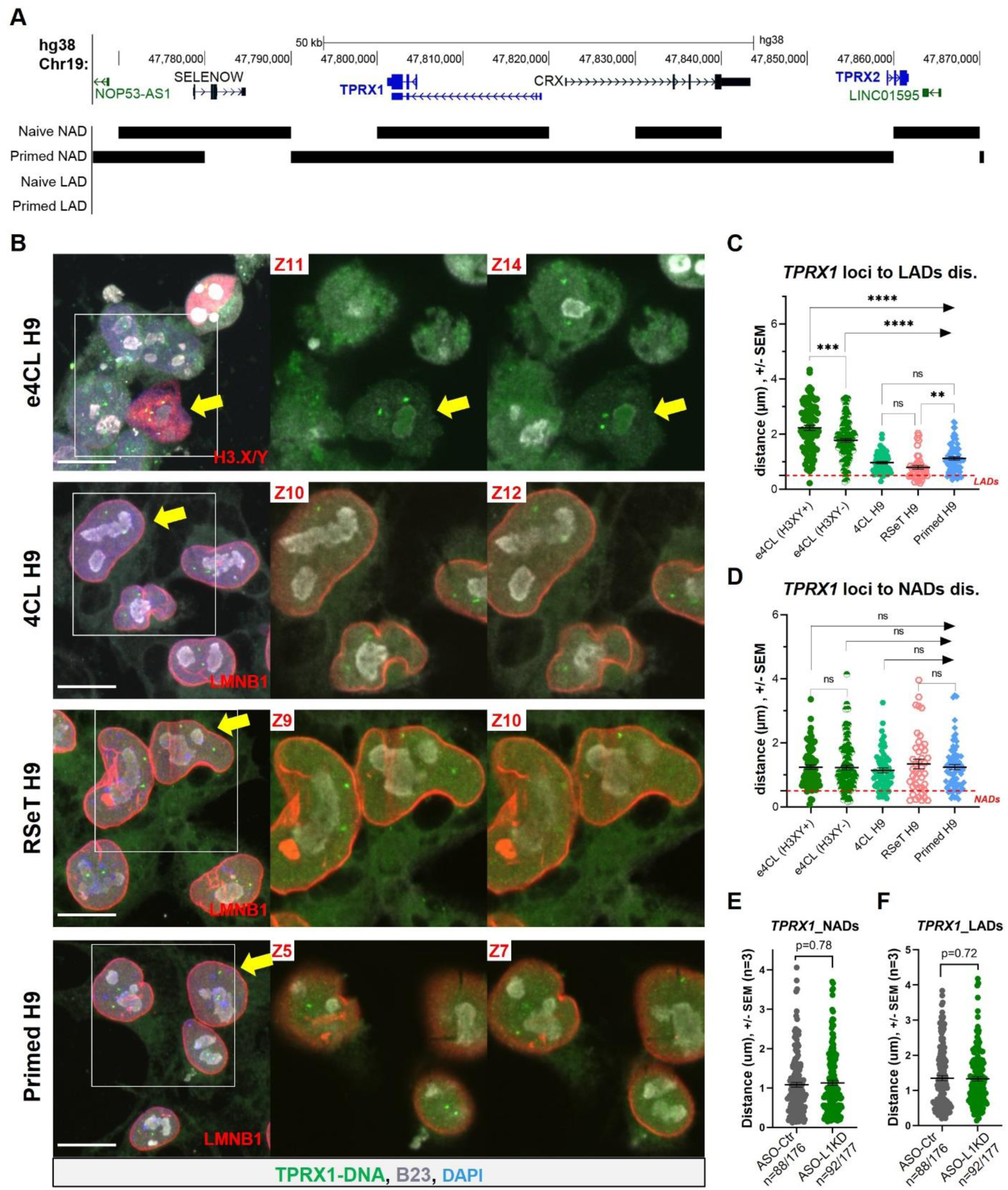
*TPRX1* loci gain association with the nucleolus and the lamina in the progression from 8CLCs to more developmentally advanced hESC states. (A) Genome browser view of the *TPRX1* region, displaying enrichment in NADs. (B) Representative image of *TPRX1* DNA-FISH and co-IF staining for B23 and LMNB1 in 4CL, RSeT and primed hESCs. In e4CL, instead of LMNB1 antibody, co-IF was carried out for H3.XY to identify 8CLCs; thus, the distance of *TPRX1* loci to the lamina in the images of e4CL cells was measured to the DAPI periphery. The boxed area in the max projected images (left panels) is enlarged and displayed as two representative z-stack (Z) images (right panels). The large yellow arrows point to representative cells exported in Videos S3-S6. Representative of two independent experiments. Scale bar, 10 µm. (C and D) Quantification of data from the *TPRX1* DNA-FISH and co-IF staining (shown in A), plotting the distance of *TPRX1* loci to (C) DAPI periphery (e4CL) or LMNB1-marked laminar domain (4CL, RSeT and primed hESCs); and (D) B23-marked nucleolus domain. If the distance is < 0.5 µm, it is defined as within LADs or NADs. Data are from two replicated independent experiments. Number of cells quantified in each group is indicated in Figure 4C. Brown-Forsythe and Welch Anova tests. ns = p > 0.05, ** = p < 0.01, *** = p< 0.001, **** = p < 0.0001. (E and F) Quantification of data from the *TPRX1* DNA-FISH and co-IF staining for B23 and LMNB1 in control and LINE1 KD RSeT+DT hESCs, plotting the distance of *TPRX1* loci to (E) nucleolar and (F) lamina domains. n=cells /*TPRX1* loci quantified. Data are from three independent experiments. Mann-Whitney test. ns = p > 0.05.

**Figure S11.**
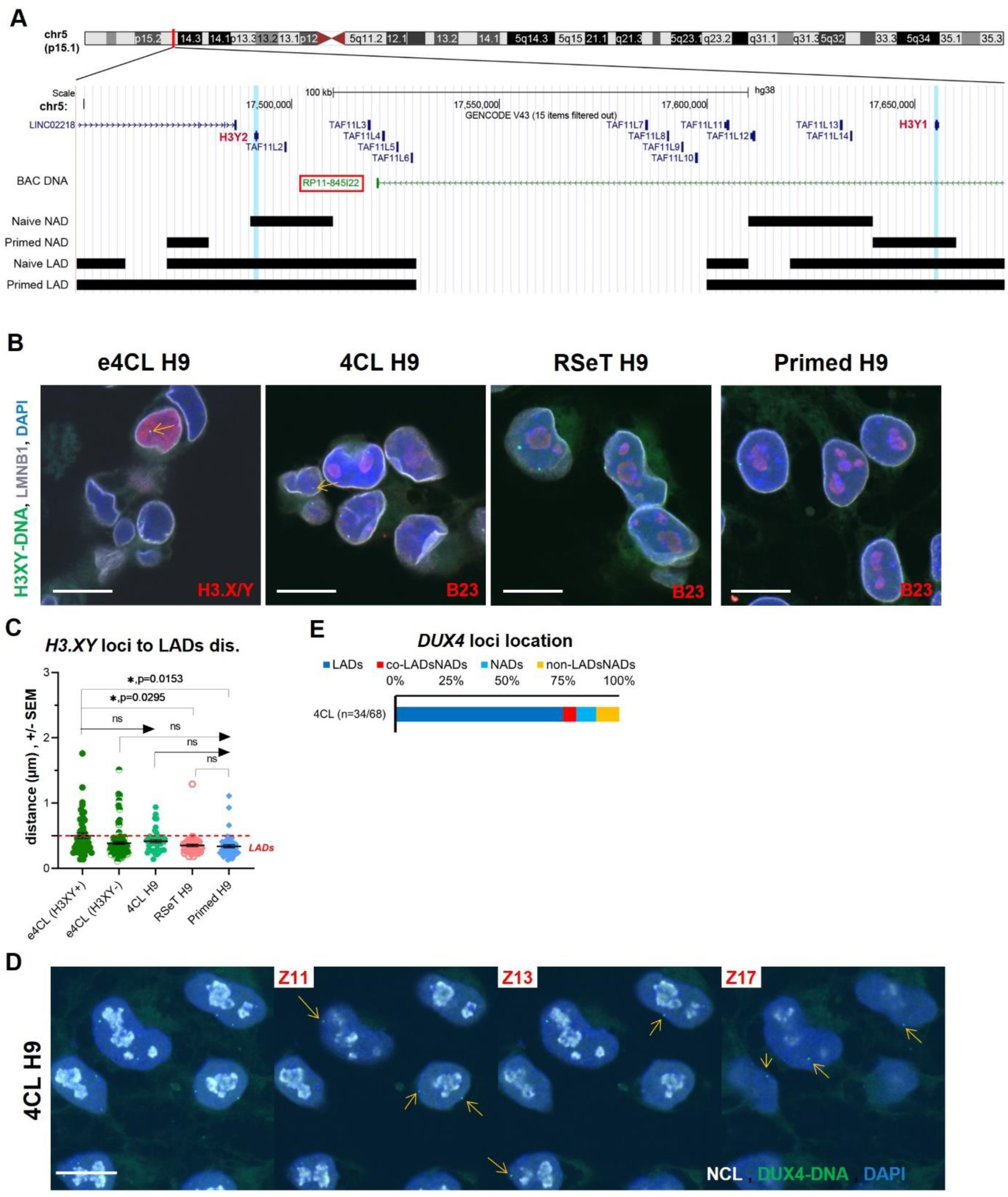
*H3.XY* and *DUX4* loci are primarily located at the lamina in hESCs. (A) Genome browser view of the *H3.XY* (annotated as *H3.Y1* and *H3.Y2*) loci, displaying enrichment in LADs and NADs. *H3.XY* DNA-FISH probes are generated with the RP11-845I22 BAC plasmid. See Table S4 for DNA-FISH probes for other loci. (B) Representative z-stack image of *H3.XY* DNA-FISH and co-IF staining for B23 and LMNB1 in 4CL, RSeT and primed hESCs. In e4CL, instead of B23 antibody, co-IF was carried out for H3.XY to identify 8CLCs; thus, the distance of *H3.XY* loci to nucleolus in the images of e4CL cells was not measured. The yellow arrows point to *H3.XY* loci in the nucleoplasm. Representative of two independent experiments. Scale bar, 10 µm. (C) Quantification of data from the *H3.XY* DNA-FISH and co-IF staining (shown in B), plotting the distance of *H3.XY* loci to the laminar domain. If the distance is < 0.5 µm, it is defined as within LADs. Number of cells quantified in each group is indicated in Figure 4D. Brown-Forsythe and Welch Anova tests. ns = p > 0.05, * = p < 0.05. Representative of two independent experiments. (D) Representative image of *DUX4* DNA-FISH and co-IF staining for NCL in 4CL hESCs. The distance of *DUX4* loci to the lamina in the images was measured to the DAPI periphery. The max projected image (left panel) is displayed as three z-stack (Z) images (right panels). The yellow arrows point to *DUX4* loci, locating frequently at the DAPI periphery. Representative of at least two independent experiments. Scale bar, 10 µm. (E) Plot of the *DUX4* location percentile in each listed category based on the distance quantification (data not shown). The definition of its location in LADs or NADs is the same as above in (C and D). n=cells / *H3.XY* loci quantified. Representative of two independent experiments.

**Figure S12.**
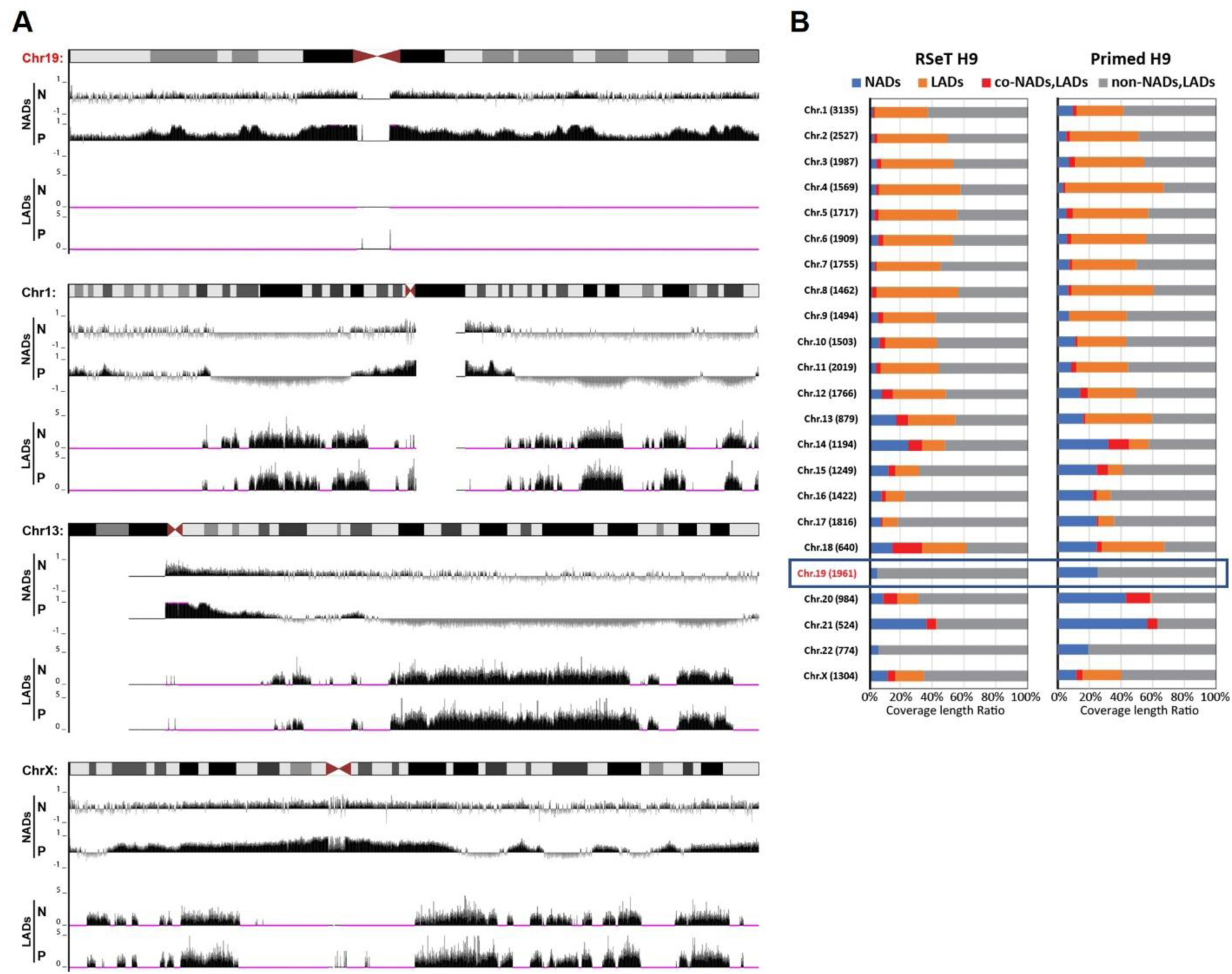
Chromosome 19 contains NADs but not LADs. (A) Representative genome browser views of LADs and NADs enrichment signatures of Chr. 19, Chr. 1, Chr. 13, and Chr. X., showing large-scale changes in NADs and very stable LADs in Naïve-like (N) and Primed (P) hESCs. (B) Plot of NADs, LADs, co-LADs/NADs, and non-LADs/non-NADs coverage as percentage to the total length in each chromosome. Chr. 19 lacks LADs in both RSeT and Primed hESCs.

**Figure S13.**
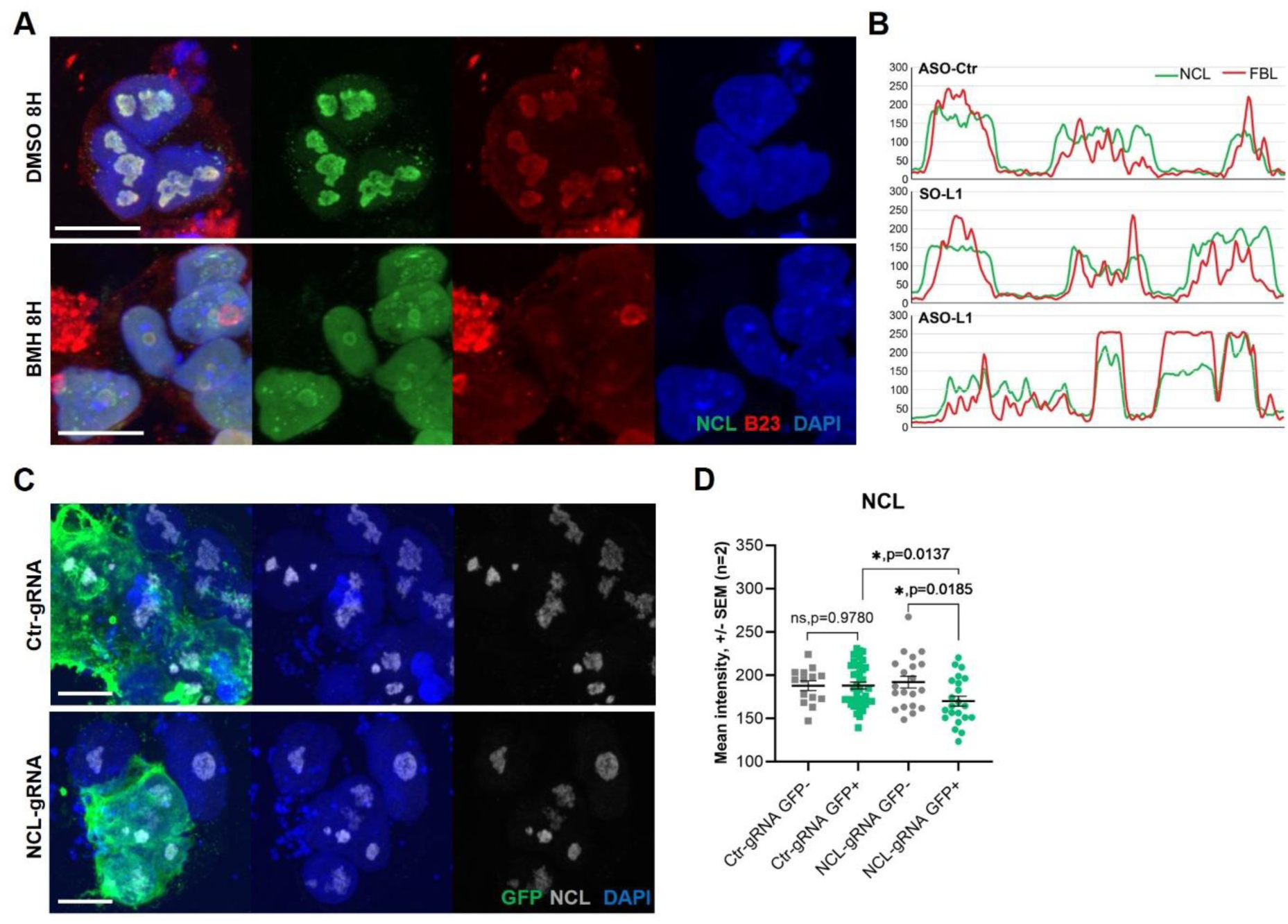
RNA Pol I inhibition and CRISPRi KD of NCL impair nucleolar architecture in hESCs. (A) NCL and B23 IF in hESCs treated with 0.25 µM BMH for 8 hours, showing disrupted nucleolar architecture, relative to DMSO controls. Representative of two independent experiments. Scale bar, 10 µm. (B) Representative NCL and FBL signal intensity profile plot across nucleolus (white-dotted line in Figure 6B), indicating an extrusion of FBL from the NCL territory in LINE1 KD cells. (C) NCL IF in dCas9-KRAB transgenic hESCs transfected with NCL-gRNAs, compared to Ctr-gRNAs as control. GFP co-IF identifies cells with positive gRNA transfection. Representative images of two independent experiments. Scale bar, 10 µm. (D) Quantification of NCL and GFP IF images (examples in C), showing significant KD of NCL protein in GFP+ transfected with NCL-gRNAs, relative to controls. Data are from two independent experiments. Welch’s t-test. ns = p > 0.05, * = p < 0.05.

**Figure S14.**
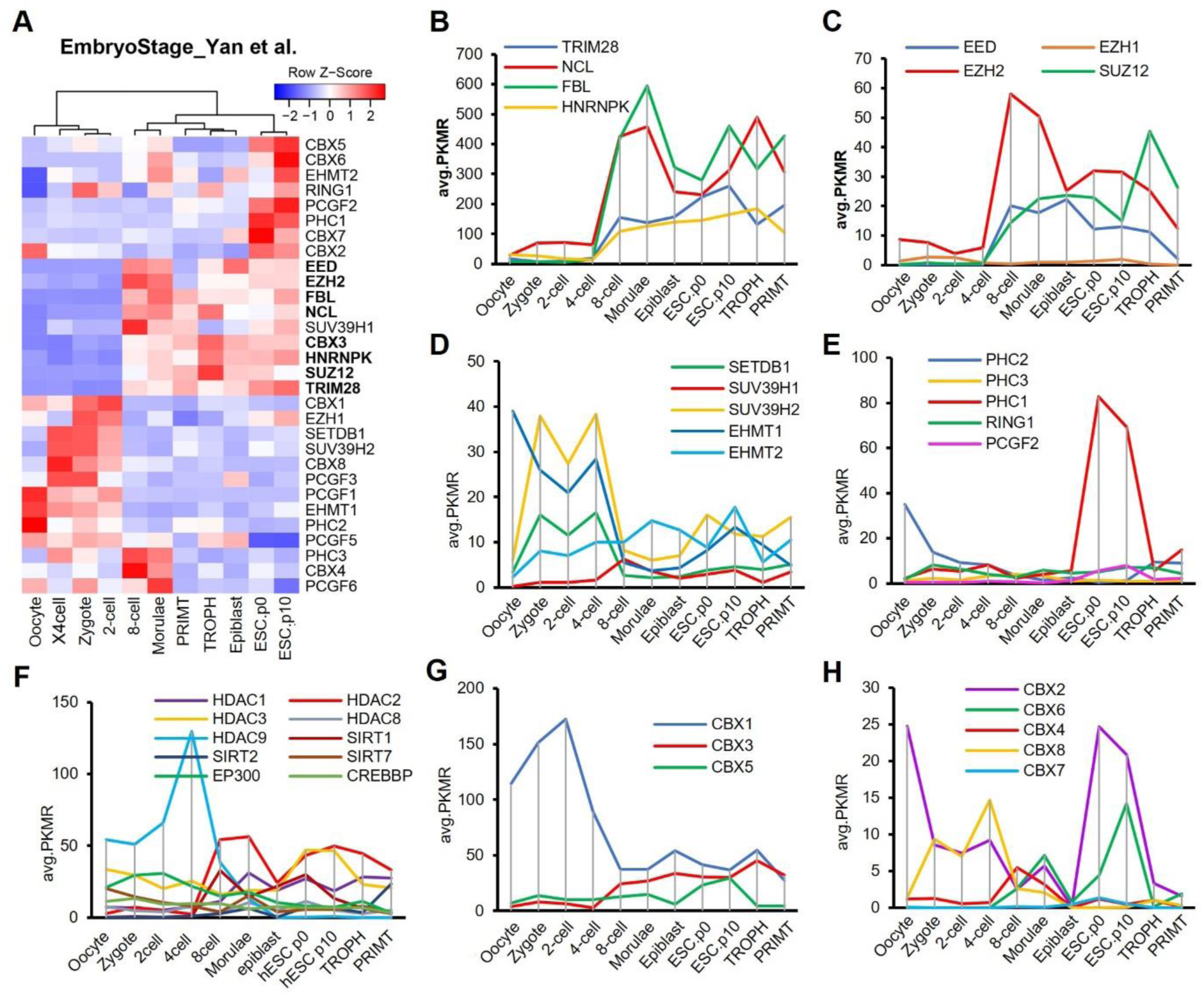
Nucleolar factors and components of PRC2 are highly induced at the 8C stage in vivo. (A) Heatmap representation of the expression of select heterochromatin and nucleolus factors during early human embryonic development, profiled using scRNA-seq^22^. Nucleolar proteins and PRC2 subunits are among genes significantly upregulated (bolded) at the 8C stage. TROPH, trophectoderm; PRIMT, Primitive endoderm. (B-H) Plot of the average expression levels (Penalized Kernel Matrix Regression, avg.PKMR of indicated genes in cells of each embryonic stage sample, profiled using scRNA-seq^22^. TROPH, trophectoderm; PRIMT, Primitive endoderm.

